# Uncovering moonlighting role of mitochondrial presequence translocase machinery in SOD1-mediated ALS pathogenesis

**DOI:** 10.64898/2026.06.03.729791

**Authors:** Tejashree Pradip Waingankar, Aakansha Paliwal, Anjali Deep, Patrick D’Silva

## Abstract

Familial Amyotrophic Lateral Sclerosis (fALS) is a fatal neurodegenerative disease, mainly caused by mutations in the superoxide dismutase 1 (SOD1) protein. Mitochondrial dysfunction is a primary hallmark of ALS pathogenesis. However, the molecular mechanism by which SOD1 mutants impair organellar health remains enigmatic. This study demonstrates that mutant SOD1 associates with the TIM23 complex in *Saccharomyces cerevisiae* via its intermembrane space (IMS) domain. In ALS-associated SOD1 mutants, both binding and expression of TIM23 complex proteins were downregulated, leading to altered translocation of the substrate protein Sdh3, a component of the electron transport chain (ETC) complex II. Disrupted Sdh3 translocation leads to mitochondrial dysfunction, evidenced by decreased ETC complex II activity, reduced functional mass, and compromised organelle integrity. Overexpression of Tim23 partially rescued mitochondrial integrity by increasing ETC complex activity and functional mass and restoring reticular morphology. Strikingly, the improved mitochondrial homeostasis in Tim23-overexpressing cells partially rescued the growth defects caused by mutant SOD1. Collectively, these findings reveal a previously unrecognized regulatory axis between mutant SOD1 and the mitochondrial pre-sequence translocase machinery, highlighting this pathway as a promising target for future ALS therapies and opening new avenues for mechanistic and translational research.

**Author Summary:** Familial Amyotrophic Lateral Sclerosis (fALS) is a progressive, fatal neuromuscular disorder marked by motor neuron degeneration. The exact cause of ALS remains unclear. Previous research links familial ALS to mutations in the superoxide dismutase 1 (*SOD1*) gene. SOD1 mutants in ALS disrupt mitochondrial protein translocation, a key mitochondrial process. The mechanism by which SOD1 mutants affect mitochondrial function and integrity by modulating presequence translocase (TIM23 complex) import is not yet understood.

The current study addresses a critical gap in ALS research by demonstrating a novel, direct interaction between SOD1 and Tim23 that regulates mitochondrial function in yeast. We found that SOD1 binds Tim23 via Tim23 IMS domain, stabilizes the Tim23^CORE^ complex, enabling Sdh3 import. Loss of SOD1, Tim23, or Tim50 destabilizes the TIM23^CORE^ complex, leading to impaired Sdh3 import and decreased ETC complex-II activity. These changes disrupt mitochondrial structure, causing fragmentation and a loss of functional mitochondrial mass in Δ*sod1*. ALS-linked SOD1 mutants show similar effects: they diminish Sdh3 import by weakening SOD1-Tim23 interaction and lowering TIM23 complex stability, resulting in punctate mitochondria and reduced mitochondrial mass. Collectively, our study identifies the SOD1-TIM23 interaction as a key regulator of mitochondrial health through Sdh3 import via the TIM23^CORE^ complex and indicates this pathway as a potential early intervention target for ALS therapy.

## Introduction

Familial Amyotrophic Lateral Sclerosis (fALS) is a progressive, fatal neurodegenerative disease and its survival is limited to 3-5 years after onset [1, 2]. ALS is marked by the selective degeneration of upper and lower motor neurons in the brain and spinal cord. However, recent reports suggest multiple cell types including astrocytes, oligodendrocytes, microglia, Schwann cells, and immune cells also contribute to disease pathogenesis [1, 3–6]. As motor neuron function deteriorates, paralysis and death due to respiratory failure occur [7]. Only two drugs, Riluzole and edaravone (Radicava), have FDA approval. At most, these increase life expectancy by a few years, but do not cure the disease [8–10]. Therefore, delineating the mechanisms underlying neuronal death is critical to identifying effective treatments for ALS.

Most ALS cases are sporadic (sALS) in nature without any identified cause associated with the disease onset and progression. However, around 10% of ALS cases occur because of the familial inheritance of the known genetic mutation (fALS). A vital cellular antioxidant, superoxide dismutase 1 (SOD1), was the first gene locus identified to be mutated in ALS, and more than 160 dominant mutations have been reported till date [11]. The pathogenesis of mutant SOD1 is not limited to fALS, but its misregulation is also detected in 5% of sALS cases, highlighting its importance in ALS development [12]. Interestingly, the SOD family has two additional members: SOD2, which is exclusively mitochondrial, and SOD3, which is secreted outside the cell. However, only mutations in SOD1 were observed to cause motor neuron death in fALS [13]. The critical function of SOD1 is to dismutase highly toxic superoxide radicals. Hence, previously, it was believed that the defects in the enzymatic activity were responsible for ALS. In contrast, many disease-causing mutations subsequently identified retained significant dismutase activity [14]. Besides, later studies also showed the accumulation of misfolded oligomers of the mutant protein as the primary cause of toxicity in SOD1-mediated ALS [15,16]. Although most of the fraction of SOD1 is localized in the cytoplasm, a small fraction (∼5%) resides within the intermembrane space (IMS) of mitochondria [17]. The function of this mitochondrial SOD1 pool remains elusive and has gained greater significance since its accumulation causes organelle dysfunction in ALS neurons [18]. The abnormal oligomerization of mutant SOD1 led to impaired mitochondrial dynamics, axonal transport, oxidative phosphorylation, apoptosis, and mitophagy [19–22]. However, the molecular mechanism by which mutant SOD1 impairs mitochondrial function remains unknown. Intriguingly, a high-throughput study proposed altered mitochondrial biogenesis in the mutant SOD1 transgenic mice model [23].

Mitochondrial biogenesis is regulated by the import machinery of the outer membrane (OM) and inner membrane (IM). The OM translocase (TOM) complex serves as the main import pore for most mitochondrial-targeted proteins [24]. The IM features two essential complexes: the translocase of IM (TIM22) and the presequence translocase machinery (TIM23) [25]. The TIM23 translocase import pre-proteins from TOM across the IM. After pre-proteins are translocated across IM, they are directed to the matrix or the IM by the import motor or the TIM23 sort complex, respectively [26–27]. Matrix import is driven by mtHsp70 through ATP hydrolysis, while Tim21 and Mgr2 coordinate the lateral insertion of polypeptides with a hydrophobic stop-transfer signal [28–31]. Tim21 interacts with electron transport chain (ETC) subunits, linking TIM23 to oxidative phosphorylation [32, 33].

Recent findings show that mutant SOD1 increases the levels of mitochondrial TOM complex proteins Tom40, Tom22, and Tom20. At the same time, the import of many mitochondrial proteins with an N-terminal signal sequence (also known as a presequence) is downregulated in ALS-associated motor neurons [23]. This preliminary evidence suggests a functional interplay between mutant SOD1 translocation into mitochondria and the import machinery governing the organelle biogenesis. However, the underlying principle behind a direct correlation between SOD1 mutant import into the mitochondrial IMS region, modulation of presequence translocase activity, and ALS progression is largely elusive.

The current study examines the correlation between SOD1 mutant import and its functional crosstalk with the TIM23 complex in the progression of ALS pathogenesis, using *Saccharomyces cerevisiae* as a model organism. Our observation highlights that the steady-state expression of Tim23 complex components gets downregulated in the mitochondria upon either deletion or import of pathological SOD1 mutants. Moreover, SOD1 directly interacts with the IMS region of the TIM23 complex. Specifically, it modulates the import of Sdh3, a component of complex II, thereby regulating its mitochondrial activity. At the same time, mitochondrial-specific functional phenotypes can be partially restored by complementing the Tim23 complex, revealing a novel mechanism by which SOD1 mutants exhibit toxicity during ALS progression.

## Results

### A novel association between the presequence translocase machinery and Sod1

Motor neurons rely on mitochondrial oxidative phosphorylation for energy. Thus, mitochondrial dysfunction from mutant SOD1 leads to neuronal death. However, the mechanism by which mutant SOD1 accumulation in the IMS region causes mitochondrial dysfunction remains unclear. The observed changes in mitochondrial protein levels were notable, as defects in the import of nuclear-encoded proteins into mitochondria can severely impair mitochondrial function and health [23].

To assess the potential link between mutant Sod1 and the TIM23 complex, the genomic copy of *SOD1* (Δ*sod1*) was deleted from the *S. cerevisiae* strain (PJ53) wild type (WT) by homologous recombination. The resulting strains were transformed with an empty vector (EV) or ySod1 to analyze growth phenotype. Deletion of *SOD1* caused a growth-sensitive phenotype at all tested temperatures. This phenotype was rescued by ySod1 complementation (**Fig 1A**, **S1A-C Fig**). Immunoblot analysis confirmed the presence of these strains using antibodies specific to the FLAG-tag on ySod1. Ydj1 was used as a cytoplasmic loading control (**Fig 1B**). To check if *SOD1* deletion affects mitochondrial IM translocase components, mitochondria were isolated from cells grown at 34℃. These were used to estimate protein steady-state levels. The expression of TIM22 complex components Tim54, Tim22, and Tim10 was comparable between WT and Δ*sod1* (**Fig 1C**). The levels of Pam18, Tim21, and Por1 remained unchanged in Δ*sod1* cells (**Fig 1C**). Notably, Tim23 and Tim50, component of the TIM23 complex, were downregulated in the Δs*od1* strain. This downregulation was restored to WT levels by ySod1 complementation (**Fig 1C-E**).

**Fig 1.**
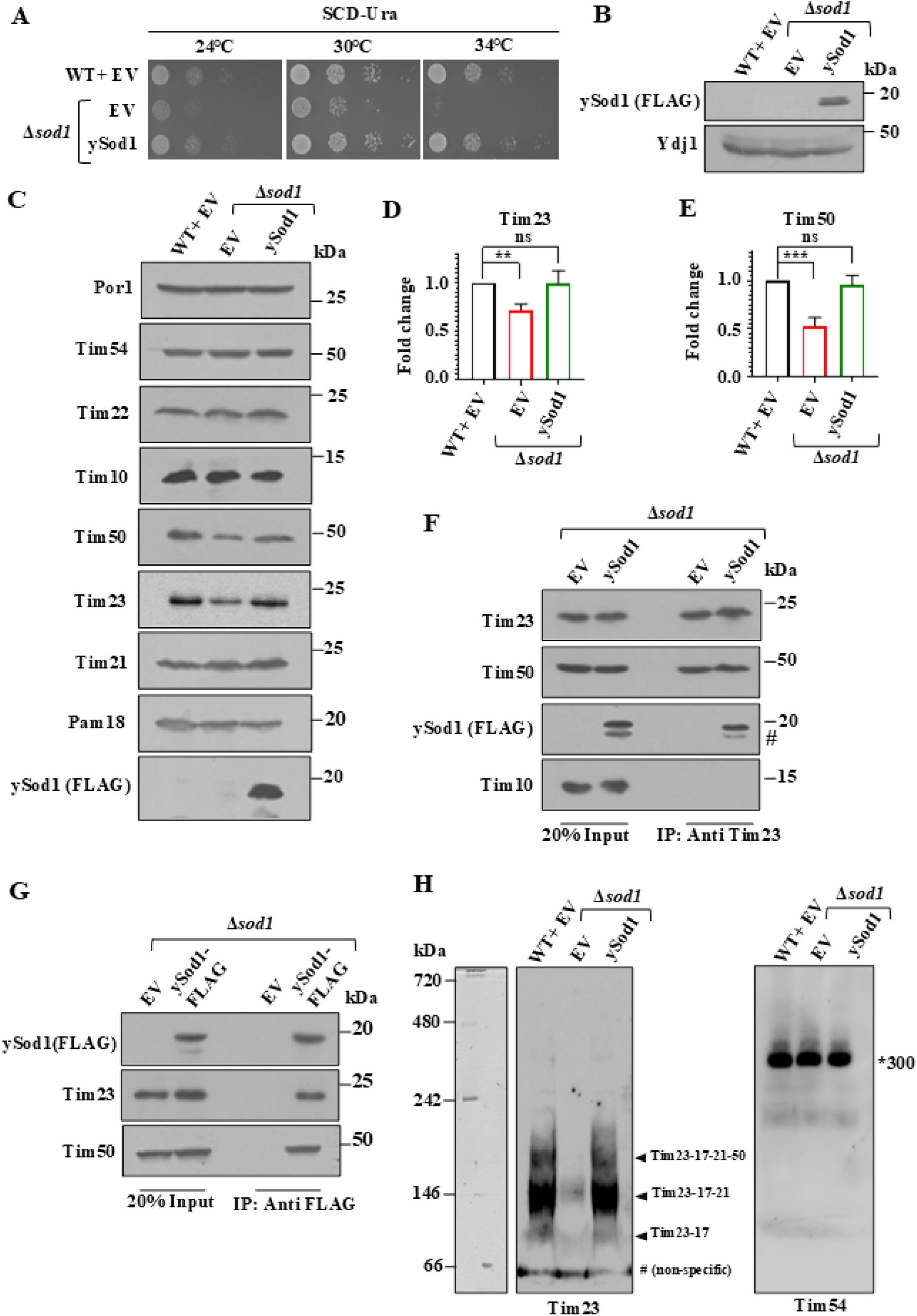
Protein expression and interaction analysis of the TIM23 complex with Sod1. **(A)** Growth complementation analysis. WT and Δ*sod1* cells were transformed with empty vector (EV) or ySod1 and selected on the SCD-Ura media. An equal number of cells was isolated from the mid-log phase and subjected to a ten-fold dilution to analyze the growth by spot assay. **(B)** Cellular expression of ySod1 levels. The cell lysate from the above-mentioned yeast strains was subjected to immunoblot analysis with the FLAG and Ydj1 antibodies. **(C)** Protein expression analysis. Yeast cells were grown at 34℃, followed by the isolation of mitochondria. An equal amount of mitochondria from the indicated strains was subjected to immunoblot analysis and detected with TIM23 complex proteins specific antibodies. **(D,E)** Quantification of protein levels. The immunoblots of the Tim23 and Tim50 were quantified using Multi Gauge software, and graphs were plotted with GraphPad Prism 8. For statistical analysis, a one-way ANOVA with Fisher’s LSD multiple comparisons test was used to compare all columns with WT+EV. Error bars indicate mean **±** standard deviation (SD) from **≥** 3 biological replicates. Asterisks indicate the *p*-value, **, *p*< 0.01. ns = not-significant. **(F,G)** co-IP analysis. Mitochondria isolated from Δ*sod1* and ySod1-FLAG strains were lysed with the non-ionic detergent digitonin. The lysed mitochondria and debris were removed, and the clear supernatant was incubated with Tim23 antibody-bound protein A Sepharose **(F)** or FLAG-linked protein G Sepharose **(G)** beads for 12h. The samples were separated by SDS-PAGE and analyzed by the protein-specific antibodies. 20% of the total mitochondrial lysate was used as the experimental input. (#) indicates a non-specific band. **(H)** Tim23-Tim50 complex stability analysis. Mitochondria isolated from WT, Δ*sod1*, and ySod1-FLAG strains were solubilized in digitonin buffer and subjected to BN-PAGE analysis, followed by immunoblotting with the indicated antibodies. Arrowheads directed indicates the three different bands which represents different forms of TIM23 complex (Tim23-17,50-21, Tim23-17-50, and Tim23-17), and the asterisk (*) indicates the TIM22 complex (∼300 kDa), taken as a control.

Tim50 is involved in the initial recognition and presorting of polypeptides exiting the TOM complex. A portion of Tim23 participates in forming the TIM-TOM super complex, which delivers precursors efficiently [34–35]. A large domain of Tim50 and Tim23 proteins is exposed in the IMS region, where SOD1 is known to translocate [36]. To gain insight into the reduced expression of Tim23 and Tim50, we hypothesized that the TIM23 complex may physically interact with Sod1. To assess the interaction between the TIM23 complex and Sod1, a co-immunoprecipitation (co-IP) analysis was performed with mitochondria isolated from the Δs*od1* and ySod1-FLAG strain. Tim50, a well-established interacting partner of Tim23, co-immunoprecipitated in both Δs*od1* and ySod1-FLAG strain (**Fig 1F**). Interestingly, ySod1 was also detected in the co-IP fraction. This suggests that Tim23 interacts with ySod1. However, this interaction was absent in Δ*sod1,* which confirmed the specificity of the co-IP (**Fig 1F**). Tim10, a component of the TIM22 complex residing in the IMS region, was used as a negative control and did not show any interaction with Tim23 (**Fig 1F**). To further confirm the binding of ySod1 to the TIM23 complex, we performed a co-IP analysis with ySod1-FLAG. Tim23 and Tim50 were observed in the co-IP fraction of ySod1-FLAG, further validating the binding of Sod1 to the mitochondrial presequence translocase machinery. Δs*od1* was used as a negative control for the experiment (**Fig 1G**).

The decrease of Tim23 and Tim50 protein level in Δ*sod1* strain, also suggests an imbalance of protein stability. So, we performed, a cycloheximide chase assay to investigate the effect of Sod1 on turn-over rate of Tim23 and Tim50. The rate of Tim23 and Tim50 degradation in the *SOD1* knockout strain was comparable to the WT and ySod1 at all analyzed time points (**S1D-F Fig**). Ydj1, a cytosolic protein with a known half-life of 11.4 hours (as listed in *Saccharomyces* Genome Database (SGD), was used as a control. Ydj1 remained stable in WT as well as Δ*sod1* strains up to 12h after cycloheximide treatment (**S1D Fig**). This result suggests that the Tim23 and Tim50 protein turnover rates remain unaltered in the absence of *SOD1*.

Tim23 and Tim50 are the essential core components of the TIM23 complex. Alterations in their expression are known to destabilize the core channel architecture [34]. Due to observed reduction in Tim23 and Tim50 expression in the Δ*sod1* strain, we subsequently evaluated the stability of the core TIM23 complex. For this, digitonin-lysed mitochondria of WT, Δ*sod1*, and ySod1 grown at 34°C were subjected to Blue-Native PAGE. Upon analysis, we observed 3 bands representing the different forms of TIM23 core complexes (Tim23-17-21-50, Tim23-17-21, and Tim23-17) in WT. However, *SOD1* deletion decreased expression of the all these core complex, which was rescued by ySod1 complementation. Additionally, TIM22 was used as the negative control in the experiment. Its levels were not affected by *SOD1* gene deletion. Collectively, this suggests that *SOD1* deletion decreases the stability of the TIM23 core complex, presumably by downregulating Tim23 and Tim50 (**Fig 1H**).

### The Tim23 IMS domain modulates Sod1-TIM23 interaction to regulate protein import

To delineate the interaction interface between Tim23/Tim50 and Sod1, we analyzed the sequences of the IMS-exposed domains of Tim23 and Tim50. The IMS domain of Tim23 (1-96 residues) is an intrinsically disordered region with a distinct binding site (71–84 residues). Tim23 substrate binding site includes more than 40 residues lying only in the IMS region [37]. The interaction between the Tim23 and Tim50 IMS domains is crucial for facilitating mitochondrial protein translocation. Mutation in L71 residue of the Tim23 IMS domain results in general import defect by altering Tim23-Tim50 interaction. For this study, we selected the L64 and L78 residues of the Tim23 IMS domain to determine the Tim23-Sod1 interaction interface [34]. Here, the L71 residue mutation was a positive control for mitochondrial protein import defect. The Tim23 residues L64, L78, and L71 were mutated from leucine to serine using site-directed mutagenesis and were analysed phenotypically on synthetic His-Trp drop-out media by spot assay and growth kinetics. The growth of *tim23_L64S_* and *tim23_L78S_* was comparable to WT at all temperatures (**S2A-E Fig**); however, *tim23_L71S_*showed temperature-sensitive phenotypes at 37°C (**S2A,E Fig**).

Mitochondrial protein translocation was investigated by subjecting whole-cell lysates from all strains to a precursor accumulation assay. In this assay, Hsp60 and Mdj1 served as model proteins, while Ydj1 was used as a loading control. The WT, Tim23_314_*, tim23_L64S_*, and *tim23_L78S_* strains did not exhibit accumulation of Hsp60 and Mdj1 precursors. In contrast, the *tim23_L71S_* strain showed accumulation of these precursors, consistent with previous findings (**S3A Fig**) [34]. To confirm that the observed import defect in *tim23_L71S_*results from an IMS domain mutation rather than altered TIM23^CORE^ protein levels, the levels of Tim23, Tim17, and Tim50 were analyzed. Protein levels were found to be comparable across all strains (**S3B Fig**). To further examine the substrate specificity of the IMS-localized Tim23 binding domain, additional analyses of Tim23-specific substrates were conducted Protein levels for ETC complexes (Nde1, Cor2, Cox4), Fe-S cluster biogenesis (Nfs1), mtDNA maintenance (Aconitase), and mitochondrial matrix chaperones (Hsp60, Mdj1) were similar in WT and Tim23 mutant strains (**S3C Fig**). In contrast, Succinate dehydrogenase 3 (Sdh3), a Complex II protein, was significantly reduced in *tim23_L78S_* and *tim23_L71S_* (**SA-C Fig**). This result suggest that L78S mutation explicitly alters specific proteins import including Sdh3 import unlike L71S exhibiting general import defect.

To get molecular insight, we evaluated Tim23 mutant’s role under *SOD1* deletion. The *SOD1* gene was knocked out in WT, Tim23_314_, and Tim23 mutant strains using homologous recombination, and the resulting strains were analyzed for growth phenotypes **(Fig 2A**, **SA-C Fig**). Interestingly, the level of Sdh3 decreased significantly in Δ*sod1* and Tim23_314_/Δ*sod1* cells compared to WT and Tim23_314_ (**Fig 2B-C**). Intriguingly, the level of Sdh3 in *tim23_L64S_*/Δ*sod1* was comparable to Tim23_314_/Δ*sod1*, which suggested that the L64 residue mutation has no further effect on Sdh3 import in the absence of *SOD1* (**Fig 2B-C**). However, in *tim23_L78S_*/Δ*sod1* and *tim23_L71S_*/Δ*sod1* cells, the level of Sdh3 showed a further decrease as compared to Δ*sod1* and Tim23_314_/Δ*sod1* (**Fig 2B-C**). The drastic decrease of Sdh3 import in *tim23_L71S_*/Δ*sod1* could be due to a general overall import defect (**S4D Fig**).

**Fig 2.**
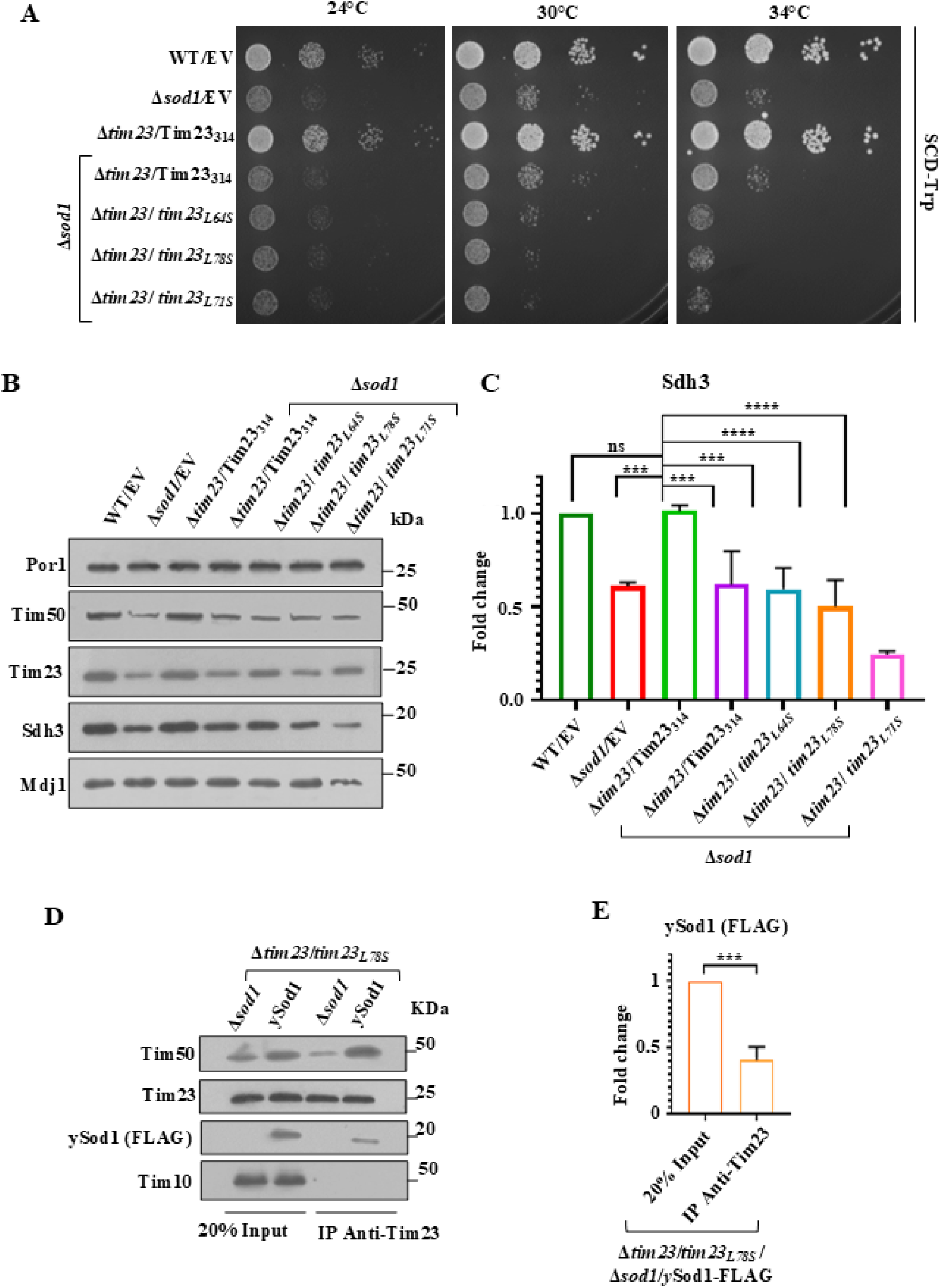
The Tim23 IMS domain modulates with Sod1-TIM23 interaction to regulate protein import. **(A)** Growth complementation analysis. WT and Tim23 mutant cells under *SOD1* knockout (Δ*sod1*) were transformed with EV and selected on SCD-Trp. An equal number of cells was isolated from the mid-log phase and subjected to a ten-fold dilution to analyze the growth by spot assay. **(B**) Protein expression analysis. Yeast cells are grown at 34°C, followed by the isolation of mitochondria. An equal amount of mitochondria from the indicated strains was subjected to immunoblot analysis, followed by the detection using protein-specific antibodies. **(C)** Quantification of protein levels. The immunoblot of Sdh3 was quantified using Multi Gauge software, and a graph was generated in GraphPad Prism 8. For statistical analysis, a one-way ANOVA with Fisher’s LSD multiple comparisons test was used to compare all columns with WT+EV. Error bars symbolize mean **±** SD from **≥** 3 biological replicates. Asterisks denote the p-value, \*\*\**p*< 0.001, \*\*\*\**p*< 0.0001. ns = not-significant. **(D)** co-IP analysis. Mitochondria isolated from the sod1 and ySod1-FLAG strains were lysed with the non-ionic detergent digitonin. The unlysed mitochondria and debris were removed, and the clear supernatant was incubated with Tim23-antibody-bound protein A sepharose beads for 12h. The samples were separated by SDS-PAGE and analyzed by the protein-specific antibodies. 20% of the mitochondrial lysate was used as the experimental input. **(E)** Quantification of protein levels. The immunoblot of Sod1 was quantified using Multi Gauge software, and a graph was generated in GraphPad Prism 8. For statistical analysis, Unpaired t-test was used to compare Input and IP-fraction column. Error bars indicate mean **±** SD from **≥** 3 biological replicates. Asterisks refers the *p*-value, **, *p*< 0.01.

Together, the interaction between Tim23 and Sod1 could be at the IMS region of the TIM23 complex, and its disruption may lead to a decrease in the import of Sdh3. To reveal the interaction interface of Sod1 and Tim23 IMS domain, *tim23_L64S_/*Δ*sod1* and *tim23_L78S_*/Δ*sod1* were transformed with EV or ySod1 and subjected to growth phenotype analysis, which confirmed that ySod1 in *tim23_L78S_*/Δ*sod1* (**S5A-D Fig**) and *tim23_L64S_*/Δ*sod1* (**S5E-H Fig**) successfully rescued the growth phenotype of *SOD1* knockout at all the tested temperatures. Mitochondria isolated from *tim23_L64S_*/Δ*sod1* and *tim23_L78S_*/Δ*sod1* transformed with EV or ySod1 strains were subjected to co-IP. Here, we observed that the level of ySod1 in the IP fraction of *tim23_L64S_*/Δ*sod1* (+ySod1) was comparable to the input fraction (**S5I-J Fig**). The results indicate that the L64S mutation in the Tim23 IMS domain may not disrupt the interaction between Sod1 and Tim23. However, a significant reduction in Sod1 expression was observed in the co-IP fraction of *tim23_L78S_*/Δ*sod1* (+ySod1) compared to the input fraction, suggesting that the L78 residue mutation could modulate Sod1 dynamic interaction with the TIM23 complex (**Fig 2D-E**).

### Tim23 IMS domain mediated Tim23-Sod1 interaction defect partially impairs mitochondrial health

To evaluate the effects of reduced Sdh3 import resulting from the defective interaction between the Tim23 intermembrane space (IMS) domain and superoxide dismutase 1 (SOD1), we assessed total mitochondrial mass and mitochondrial membrane potential, which are indicators of mitochondrial integrity. Total mitochondrial mass was measured using the cardiolipin-specific dye 10-N-nonyl acridine orange (NAO) via flow cytometry. The Δ*sod1* strain (*SOD1* deletion), as well as the *tim23_L64S_*/Δ*sod1*, *tim23_L78S_*/Δ*sod1*, and *tim23_L71S_*/Δ*sod1* double mutants (Tim23 mutants in a *SOD1* deletion background), displayed total mitochondrial masses comparable to the wild-type strain containing empty vector and Tim23_314_ (**Fig 3A**).

**Fig 3.**
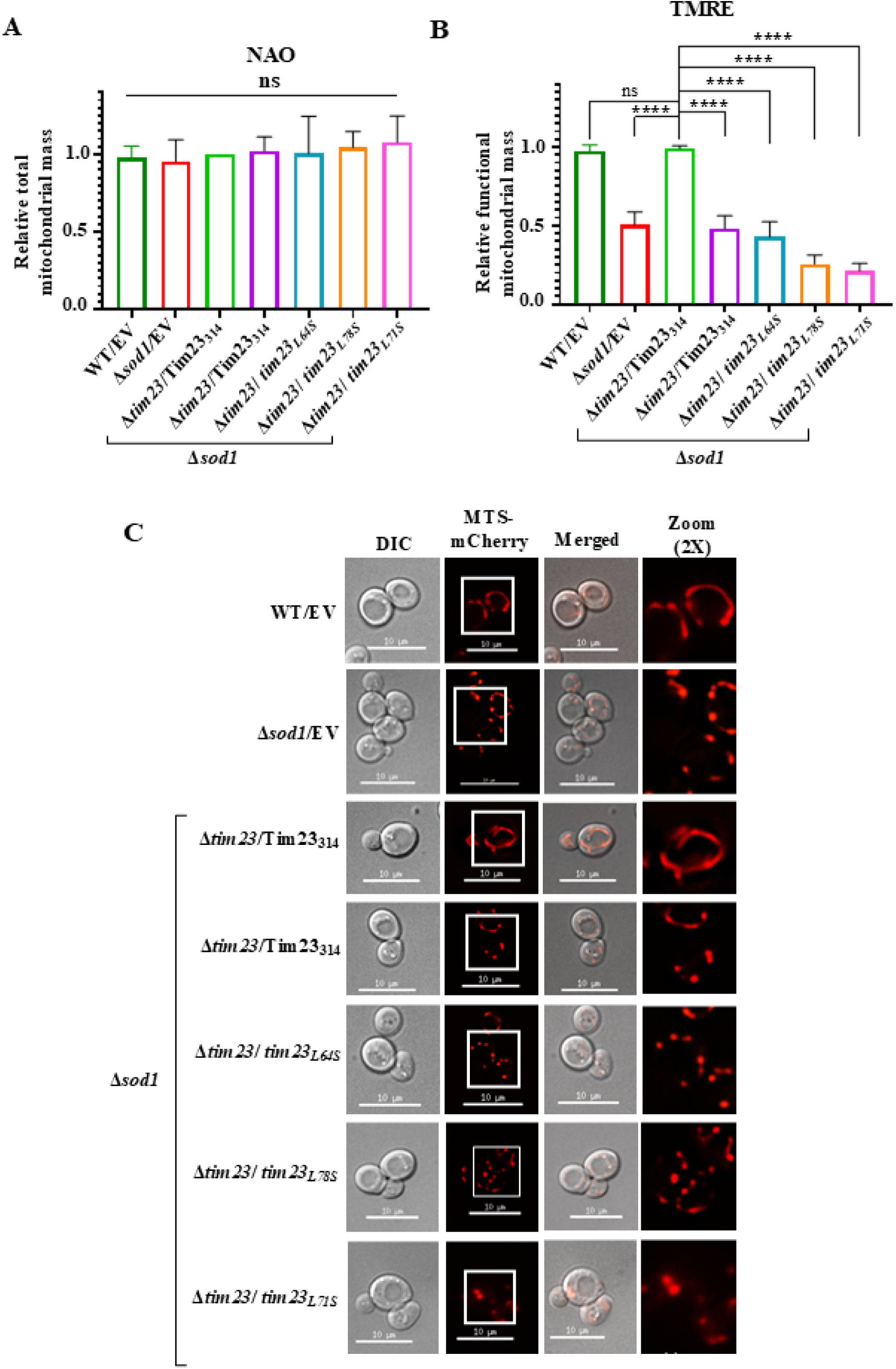
Tim23 IMS domain-Sod1 interaction defect partially impairs mitochondrial health. **(A)** Total mitochondrial mass estimation. An equal number of cells from the mid-log phase were incubated with 10 µM NAO. The cells were analyzed by flow cytometry and represented a fold change compared to Tim23_314_. **(B)** Functional mitochondrial mass assessment. Yeast cells from the above-mentioned strains were treated with 8.75 µM TMRE for 20 min at 30℃, followed by flow cytometry analysis. For statistical analysis, a one-way ANOVA with Fisher’s LSD multiple comparisons test was used to compare all columns with *Δtim23*/Tim23_314_. Error bars represent mean **±** SD from **≥** 3 biological replicates. Asterisks indicate the p-value, \*\*\**p*< 0.001, \*\*\*\**p*< 0.0001. ns = not-significant. **(C)** Mitochondrial morphology determination. Yeast cells expressing MTS-mCherry were analyzed by fluorescence microscopy to assess mitochondrial morphology. Scale bar: 10 µm.

Mitochondrial membrane potential (IM potential) was analyzed using the positively charged dye Tetramethylrhodamine methyl ester (TMRE) through flow cytometry. TMRE accumulates in active mitochondria due to their negative charge, but loss of IM potential results in TMRE escaping and reduced fluorescence. Δ*sod1* cells exhibited a significant reduction in IM potential compared to wild-type cells carrying an empty vector and Tim23_314_ cells. *Tim23_L64S_*/Δ*sod1* and *tim23_L78S_*/Δ*sod1* mutant cells also showed decreased IM potential similar to Δ*sod1* cells (**Fig 3B**). Notably, the t*im23_L71S_*/Δ*sod1* mutant demonstrated a drastic reduction in mitochondrial membrane potential, suggesting the critical role of Sod1-Tim23 IMS domain interaction in preserving mitochondrial integrity (**Fig 3B**). Additionally, mitochondrial morphology was evaluated using fluorescence microscopy as another parameter of mitochondrial integrity. Yeast strains were transformed with the MTS-mCherry vector to visualize mitochondrial networks, with the mitochondrial targeting signal (MTS) derived from subunit 9 of ATP synthase (pSU9) fused to mCherry for precise mitochondrial localization [38]. The mitochondrial morphology exists in different form including tubular, reticular, and fragmented. (**S6A Fig**). The wild type cells exhibited reticular mitochondrial morphology, while Δ*sod1* cells displayed fragmented mitochondria indicative of mitochondrial dysfunction. Tim23/Δ*sod1* cells similarly showed fragmented mitochondria. Mutations in the Tim23 IMS domain in strains *tim23_L64S_*/Δ*sod1*, *tim23_L78S_/*Δ*sod1*, and t*im23_L71S_/*Δ*sod1* further increased the percentage of cells with fragmented mitochondria (**Fig 3C, S6B Fig**).

These results suggested that loss of Sod1 disturbs mitochondrial integrity. This effect is further exacerbated by the Tim23 IMS domain mutation. Together, the Tim23 IMS domain and Sod1 interaction could play a crucial role in maintaining mitochondrial integrity.

### ALS-causing hSOD1 mutants exhibit cytotoxicity in yeast cells

The architecture of the TIM23 complex and its overall import mechanism have remained conserved during evolution. Therefore, we analyzed the interaction between SOD1 and the TIM23 complex in humans. Yeast and human SOD1 (hSOD1) share significant sequence identity (about 60%) (**Fig 4A**). We isolated mitochondria from HEK293T cells and performed co-IP using Tim23 antibody-bound protein A Sepharose beads. The co-IP fraction detected Tim50 and Tim17, both known components of the human TIM23 complex (**Fig 4B**). Notably, hSOD1 was immunoprecipitated with Tim23. This suggests that the interaction between SOD1 and the TIM23 complex is evolutionarily conserved (**Fig 4B**). We used beads only and Tom70, a component of the OM, TOM complex, as negative controls (**Fig 4B**).

**Fig 4.**
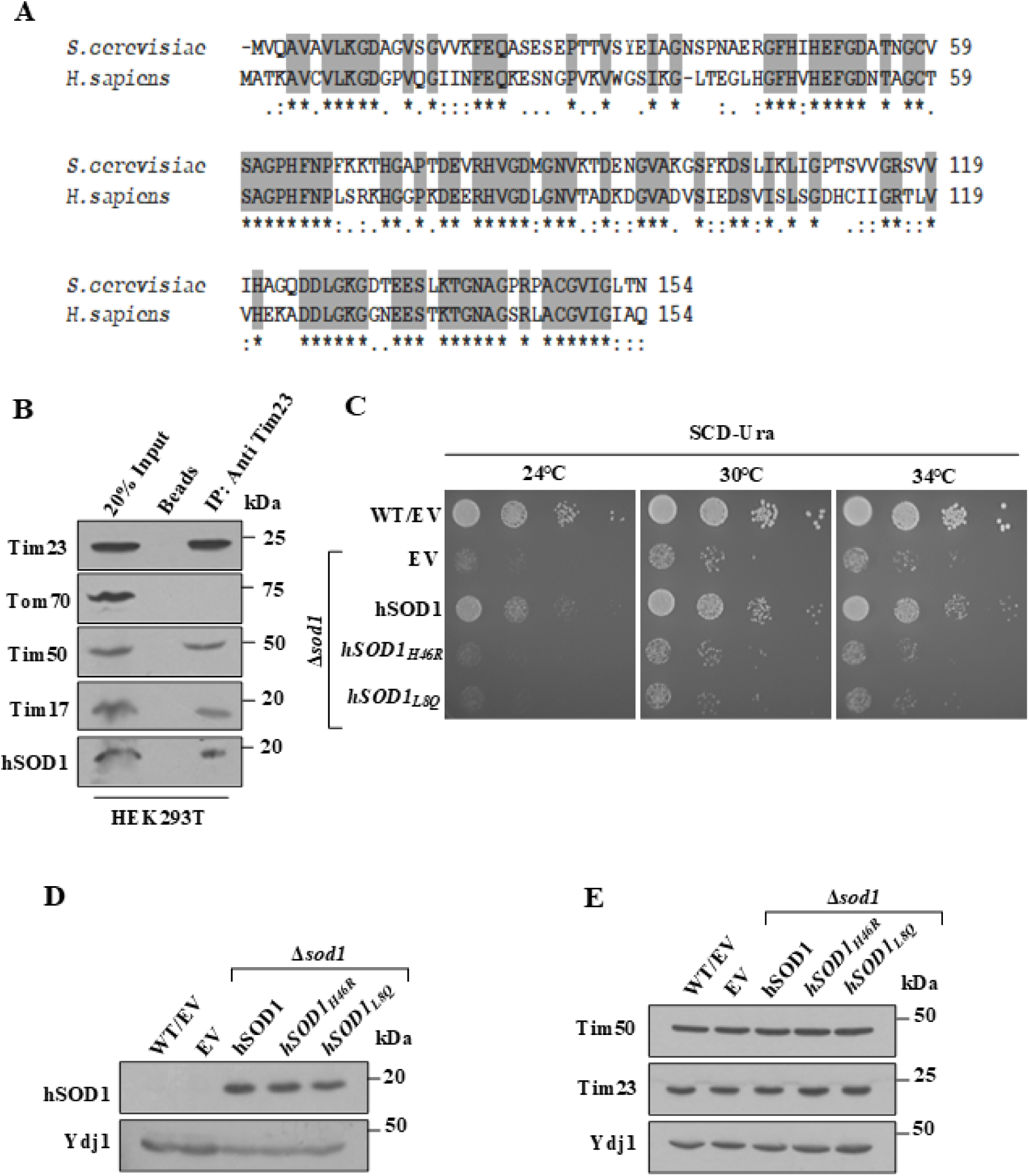
Growth phenotype analysis of ALS-linked hSOD1 mutants. **(A)** Multiple sequence alignment. Yeast and human SOD1 were analyzed for sequence similarity using Clustal Omega. Identical residues are marked in grey. **(B)** co-IP assay of human mitochondria. Mitochondria isolated from HEK293T cells were subjected to a co-IP assay with Tim23 antibody-bound protein A Sepharose beads. The co-IP fractions were separated by SDS-PAGE and analyzed by Western blot with antibodies specific against human proteins. The input for the experiment was 20% mitochondrial lysate. **(C)** Spot assay for the hSOD1 mutants. Equal amounts of hSOD1 and the mutants were isolated from mid-log phase, spotted onto SCD-Ura, and incubated at the indicated temperatures. Images were obtained after 36h. **(D)** Cellular expression of hSOD1. Cell lysates from the indicated strains were separated by SDS-PAGE and detected using protein-specific antibodies. **(E)** Cellular expression of TIM23 complex proteins: Tim23 and Tim50 in ALS-linked hSOD1 mutant strains at 30°C. Cell lysates from the indicated strains were separated by SDS-PAGE and detected using protein-specific antibodies. Ydj1 was used as a loading control.

Since the association of SOD1 with the presequence translocase machinery is evolutionarily conserved, the significance of ALS-linked hSOD1 mutants on mitochondrial biogenesis was evaluated. Two disease-causing mutants of hSOD1, *hSOD1_H46R_,* with reduced enzymatic function, and *hSOD1_L8Q_,* having significant dismutase activity, were selected to assess the dismutase-independent consequences on mitochondrial biogenesis [39–40]). To determine the cellular growth pattern in hSOD1 mutants, we expressed them in yeast cells lacking a genomic SOD1 copy and analyzed growth phenotypes. Upon spot assay, the Δ*sod1* cells displayed a temperature-sensitive phenotype, which was complemented by the expression of hSOD1 (**Fig 4C**, **S7A-C Fig**). Interestingly, both the mutants of hSOD1 showed growth defects at all the temperatures tested, underlining the cytotoxic nature of the mutants (**Fig 4C, S7A-C Fig**). The strains were confirmed by immunoblot analysis using an hSOD1-specific antibody and Ydj1 as a cytoplasmic loading control (**Fig 4D**). We also analyzed the steady-state expression of Tim23 and Tim50 in cell lysates prepared at 30℃. The levels of the TIM23 complex components remained unaltered in mutants compared to hSOD1. Altogether, overexpression of ALS-linked SOD1 exerted growth defect, however cellular Tim23 and Tim50 level remains unaltered at 30°C (**Fig 4E**)

### Cross-talk between hSOD1 mutants and the TIM23 complex modulates cellular health

To further evaluate the role of the presequence translocase machinery in hSOD1 mutants with growth defects, we analyzed the steady-state expression of Tim23 and Tim50. Mitochondria were isolated from cells grown at 34℃ and analyzed by immunoblotting. Intriguingly, Tim23 and Tim50 were downregulated in *hSOD1_H46R_*and *hSOD1_L8Q_* cells compared to hSOD1 (**Fig 5A-C**). Tim22, an IM constituent of the TIM22 complex, was used as a loading control (**Fig 5A**).

**Fig 5.**
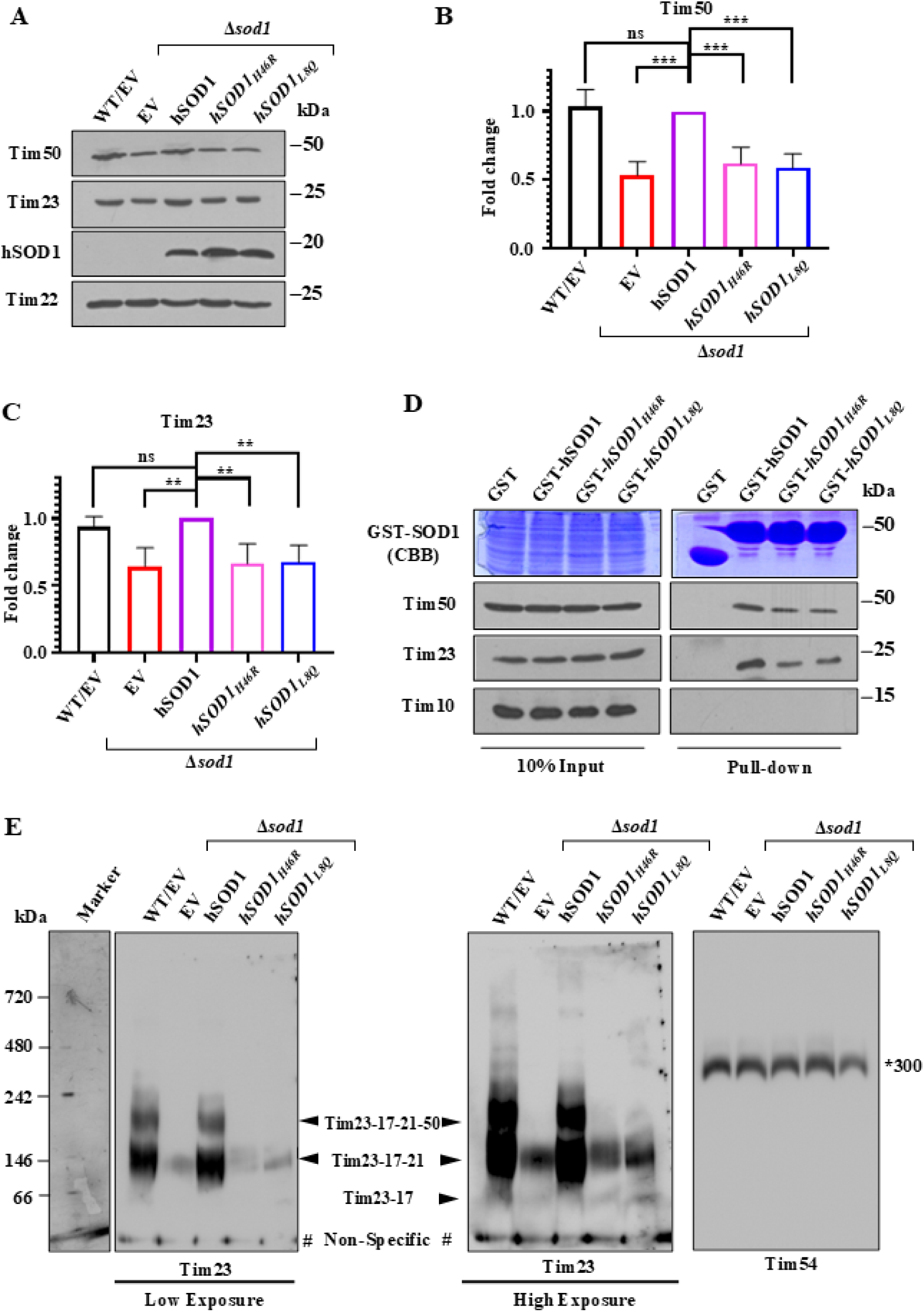
The correlation between the TIM23 complex stability in hSOD1 mutants. **(A)** Mitochondrial expression of TIM23 complex proteins. Mitochondria isolated at 34℃ were used to determine the steady-state levels of the TIM23 complex proteins. **(B,C)** Quantification is based on three independent experiments with statistical analysis performed using one-way ANOVA with Fisher’s LSD multiple comparisons test used by comparing all the columns with the hSOD1. The error bar indicates mean **±** SD for **≥** 3 different biological replicates. The asterisks indicate p values, ***, *p*< 0.001; **, *p*< 0.01. **(D)** GST pull-down assay. An equal amount of WT mitochondria was lysed with 0.5% NP-40, and the clear lysate after centrifugation was incubated with an equimolar concentration of GST-tagged hSOD1 or mutants. The samples were separated by SDS-PAGE and analyzed by immunoblot assay. Equal loading is represented by Coomassie Brilliant Blue (CBB)-stained gels. **(E)** Tim23-Tim50 complex stability analysis. Mitochondria isolated from WT, Δ*sod1*, hSOD1, *hSOD1_H46R_* and *hSOD1_L8Q_*strains were solubilized in digitonin buffer and subjected to BN-PAGE analysis, followed by immunoblotting with the indicated antibodies. Black arrowheads indicate 3 bands of different forms of TIM23 complex [Tim23-17-50-21, Tim23-17-50, and Tim23-17 (high exposure)] and the asterisk (*) indicates the TIM22 complex (∼300 kDa), taken as a loading control. Here, # symbol indicates non-specific band.

Reduced expression of TIM23 complex components in hSOD1 mutants suggested plausible alterations in their association. We tagged hSOD1 and its mutants with GST and performed a pull-down assay. An equal amount of GST alone and GST-tagged hSOD1 were incubated with mitochondrial lysate from the WT yeast strain. The hSOD1 pull-down fraction detected Tim23 and Tim50, confirming their evolutionarily conserved interaction (**Fig 5D**). *hSOD1_H46R_* and *hSOD1_L8Q_*, however, showed decreased interaction with Tim23 and Tim50 (**Fig 5D**). GST alone did not bind any tested proteins. Tim10, a part of the TIM22 complex, served as a negative control and was not detected in the pull-down fraction (**Fig 5D**).

Because Tim23 and Tim50 expression are reduced in the ALS-linked SOD1 mutant strain, we next evaluated the stability of the core TIM23 complex. To do so, we lysed mitochondria from WT, Δ*sod1*, hSOD1, *hSOD1_H46R_,* and *hSOD1_L8Q_* grown at 34°C with digitonin and subjected them to Blue-Native PAGE assay. Analysis showed three major bands of the TIM23^CORE^ complex [Tim23-17-21-50, Tim23-17-21, and Tim23-17(High Exposure)] in WT and hSOD1 strains. In contrast, ALS-linked SOD1 mutants decreased the expression of these core complexes. Furthermore, TIM22 served as a control, and its expression remained unaffected despite ALS-linked SOD1 mutant overexpression. Together, these findings suggest that ALS-linked SOD1 mutants compromises the TIM23^CORE^ complex’s stability, likely by downregulating Tim23 and Tim50 (**Fig 5E**).

The decreased expression and interaction of Tim50 and Tim23 in hSOD1 mutants pointed to a novel mechanism. ALS-associated mutants may regulate mitochondrial biogenesis this way. To test if overexpression of TIM23 complex components can rescue hSOD1 mutant growth defects, we expressed the core channel-forming unit Tim23 under the *TEF* promoter in the pRS414 yeast expression vector.

The WT or the hSOD1 strain showed comparable growth between EV control and Tim23 overexpressed conditions (**Fig 6A**, **S8A-C Fig**). However, interestingly, Tim23 overexpression partially rescued the growth defects exerted by the mutant hSOD1 at all the temperatures tested (**Fig 6A**, **S8A-C Fig**). The overexpression of Tim23 was confirmed by the immunoblot analysis in the cell lysate isolated from yeast strains, and Ydj1 was used as the loading control (**Fig 6B**). To further ascertain that the rescue of the growth phenotype is only because of Tim23, another mitochondrial IM protein, Dicarboxylate carrier 1 (Dic1), was overexpressed. Dic1 is a metabolite carrier protein imported by the TIM22 complex. The WT or hSOD1 strains did not show any alterations in growth pattern upon Dic1 overexpression. Moreover, Dic1 overexpression did not rescue the growth defects in mutant hSOD1 (**Fig 6C**, **S8D-F Fig**). The protein-specific antibody detected Dic1 and Ydj1 levels, demonstrating equal protein loading in the experiment (**Fig 6D**). Together, the cellular growth defects caused by the mutant hSOD1 can be partially rescued by the overexpression of the presequence translocase machinery protein, Tim23.

**Fig 6.**
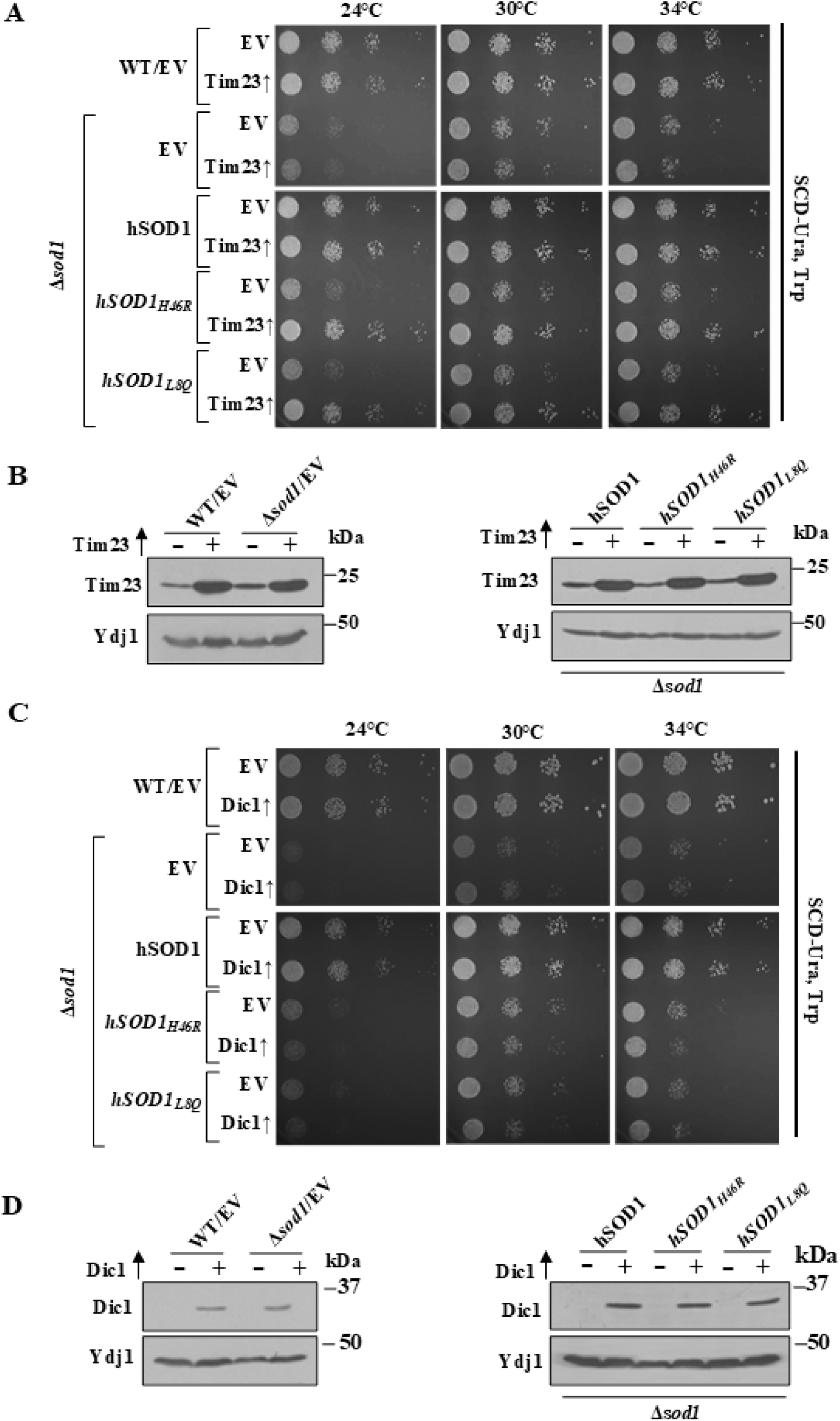
Growth assessment of Tim23 or Dic1 overexpression in hSOD1 mutants. **(A)** Temperature sensitivity assay. Yeast strains expressing an EV or a Tim23 overexpression plasmid were subjected to growth phenotype analysis at the above-mentioned temperatures. The images were acquired 48h after the growth. **(B)** Detection of protein overexpression. Cell lysates from the indicated strains were separated by SDS-PAGE and analyzed by immunoblotting with antibodies specific for Tim23 and Ydj1. **(C)** Spot analysis of Dic1 overexpressed strains. Yeast strains expressing an EV or a Dic1 overexpression plasmid were grown to mid-log phase, then diluted and spotted onto SCD-Ura and Trp minimal media. Cells were incubated at the indicated temperatures, and images were taken for representation after 48h. **(D)** Steady-state expression assessment. The cell lysate was prepared from the indicated yeast strains, and an equal amount of protein was separated by SDS-PAGE, followed by detection using Dic1 and Ydj1-specific antibodies.

### The interaction between the TIM23 complex and hSOD1 is critical for maintaining ETC complex activity

The primary function of the presequence translocase machinery is the import of polypeptides into the matrix and IM of mitochondria. Therefore, the substrates of the TIM23 complex were identified to estimate possible targets regulated by hSOD1 binding.

Mitochondria isolated at 34℃, were used to evaluate the steady-state levels of TIM23 complex substrates involved in various mitochondrial functions. The matrix chaperones, Hsp60 and Mdj1, remained unaltered between hSOD1 and the mutants (**Fig 7A**). Similarly, the TIM22 complex components, imported by the presequence translocase machinery, were found to be comparable in the case of hSOD1 and mutants. Additionally, Aconitase, required for the mtDNA maintenance, and Nfs1, essential for the biogenesis of the Fe-S cluster, remained unchanged upon mutation of hSOD1 (**Fig 7A**). Since the presequence translocase machinery is directly linked to the ETC complex by Tim21, we assessed the levels of ETC complex proteins. The Cox4 (complex IV), Cor2 (Complex III), and Nde1 (component of Complex I in yeast) expression was observed to be similar in hSOD1 mutants to that of WT hSOD1.

**Fig 7.**
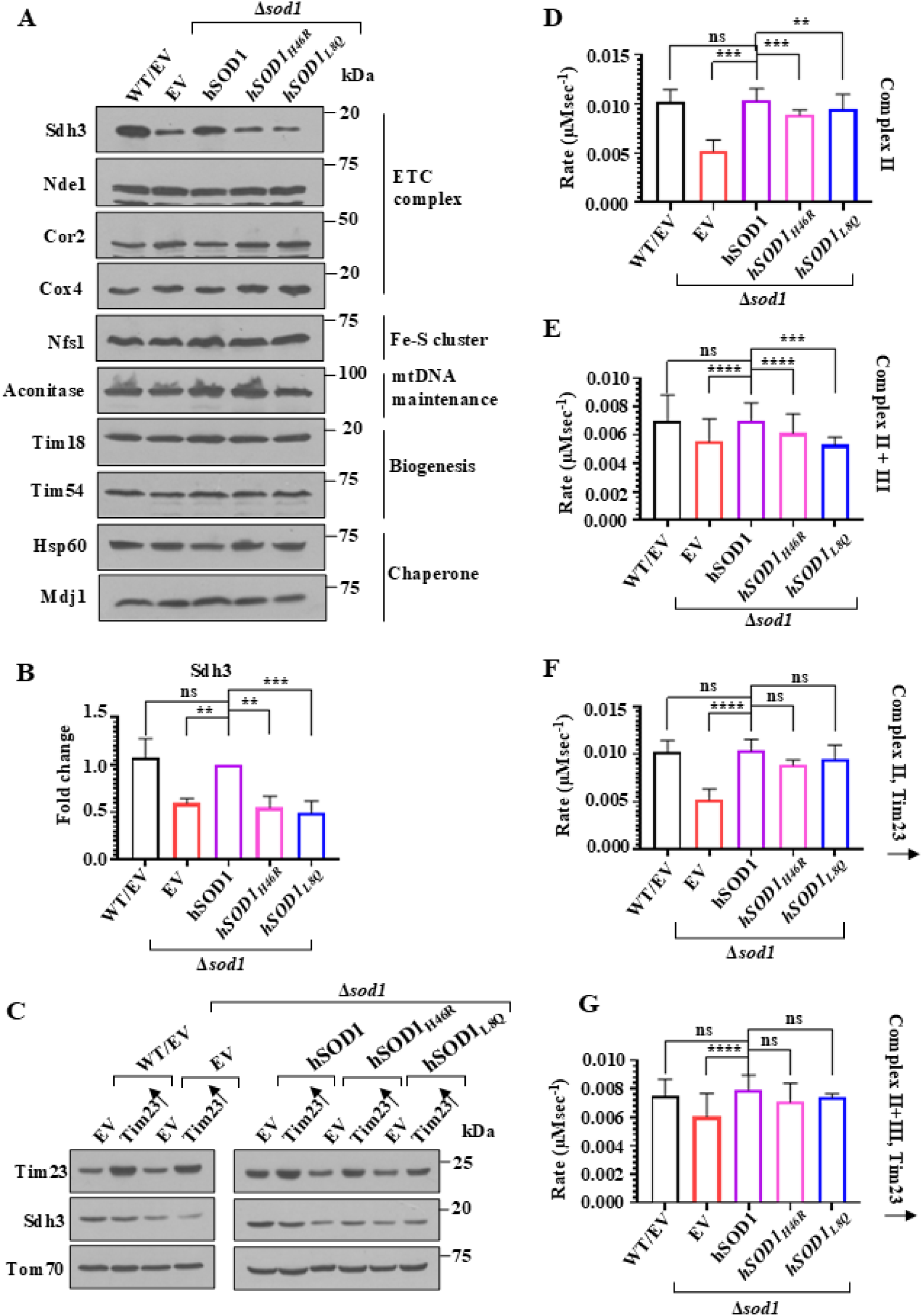
Tim23 overexpression partially rescues ETC complex activity. (A,B) Analysis of TIM23 complex substrates. The mitochondria isolated from the indicated yeast strains at 34℃ were evaluated for TIM23 complex substrates, which perform diverse functions. The immunoblots obtained from three different experiments were quantified and analyzed by one-way ANOVA Fisher’s LSD multiple comparisons test for the significance. The asterisks denote p values, ****, *p*< 0.0001; **, *p*< 0.01). **(B)** Sdh3 expression assessment. An equal amount of mitochondria isolated from EV or Tim23 overexpression strains was evaluated for the steady-state expression of the indicated proteins. **(D-G**) ETC complex activity. 30 µg of isolated mitochondria from the indicated yeast strains were incubated with succinate to activate Complex II. The resultant activity was assayed by the decrease in absorbance of DCPIP at 600 nm for Complex II **(D, F)** and an increase in Cytochrome C absorbance at 550 nm for Complex II+III **(E, G).** Error bars represent mean **±** SD from **≥** 3 biological replicates. Asterisks indicate the *p*-value, \*\**p*<0.01, \*\*\**p*< 0.001, \*\*\*\**p*< 0.0001. ns = not-significant.

However, succinate dehydrogenase 3 (Sdh3, Complex II) expression was notably decreased in Δ*sod1* cells. Complementation with hSOD1 restored Sdh3 levels. Strikingly, *hSOD1_H46R_* and *hSOD1_L8Q_* showed a significant decrease (∼50%) in the Sdh3 expression compared to hSOD1 (**Fig 7A-B**). Additionally, we determined Sdh3 levels in mitochondria isolated from Tim23-overexpressing strains. The overexpression of Tim23 partially restored the Sdh3 expression in hSOD1 mutants (**Fig 7C**). Sdh3 has a dual localization at the mitochondrial IM, wherein, apart from Complex II, it interacts with Tim18 to provide structural stability to the TIM22 complex [41]. The function of Sdh3 at the TIM22 complex is always in correlation with Tim18, but the expression of Tim18 was not altered in hSOD1 mutants, suggesting that the interaction of the TIM23 complex and hSOD1 might regulate the ETC complex activity via complex II.

To validate this possibility, the ETC complex II and the combined activity of complexes II and III were tested. Upon analysis, complex II activity was decreased in Δ*sod1,* which was restored by the expression of hSOD1 (**Fig 7D**). Remarkably, *hSOD1_H46R_ a*nd *hSOD1_L8Q_* resulted in a more than 50% decrease in the Complex II activity, in correlation with the reduced Sdh3 protein levels (**Fig 7D**). Intriguingly, the combined function of Complex II+III was also downregulated in Δ*sod1* and mutant hSOD1, suggesting the overall decrease in ETC complex activity upon expression of hSOD1 mutants (**Fig 7E**). To study the effects of Tim23 overexpression on hSOD1 mutants induced ETC complex defects, the Complex II and Complex II+III activities were examined from the mitochondria isolated from the Tim23 overexpressed strains at 34℃. The overexpression of Tim23 in Δ*sod1* showed a moderate decrease in Complex II and Complex II+III activities compared to hSOD1 (**Fig 7F-G**). Interestingly, Tim23 overexpression in *hSOD1_H46R_* and *hSOD1_L8Q_* resulted in partial rescue of Complex II and Complex II+III activity compared with hSOD1 (Fig 7F-G). Together, the overexpression of Tim23 can partially rescue the defects in ETC complex activity caused by ALS-linked hSOD1 mutants.

### Tim23 overexpression partially restores mitochondrial integrity in hSOD1 mutants

The direct consequences of ETC complex activity are seen in the mitochondrial inner membrane (IM) potential and its dynamics. These factors can be correlated to organellar dysfunction. Therefore, the mitochondrial IM potential was determined by flow cytometry analysis. The cardiolipin-specific NAO dye was used to evaluate total mitochondrial mass. The Δ*sod1* cells showed a mitochondrial mass comparable to that of WT and hSOD1 (**Fig 8A**). In contrast, hSOD1 mutants showed a significant increase in total mitochondrial mass compared to hSOD1 (**Fig 8A**). Inner membrane potential was estimated by staining yeast cells with TMRE, followed by flow cytometry. TMRE is a positively charged dye that accumulates in active mitochondria due to their negative membrane potential. Loss of inner membrane potential causes TMRE to escape, reducing fluorescence. The Δ*sod1* group showed a drastic decrease in inner membrane potential. Mutant hSOD1 showed more than a 50% reduction in mitochondrial inner membrane potential compared to hSOD1 (**Fig 8B**). The slight increase in total mitochondrial mass, together with a severe decrease in functional mitochondria, pointed to organelle dysfunction in hSOD1 mutants.

**Fig 8.**
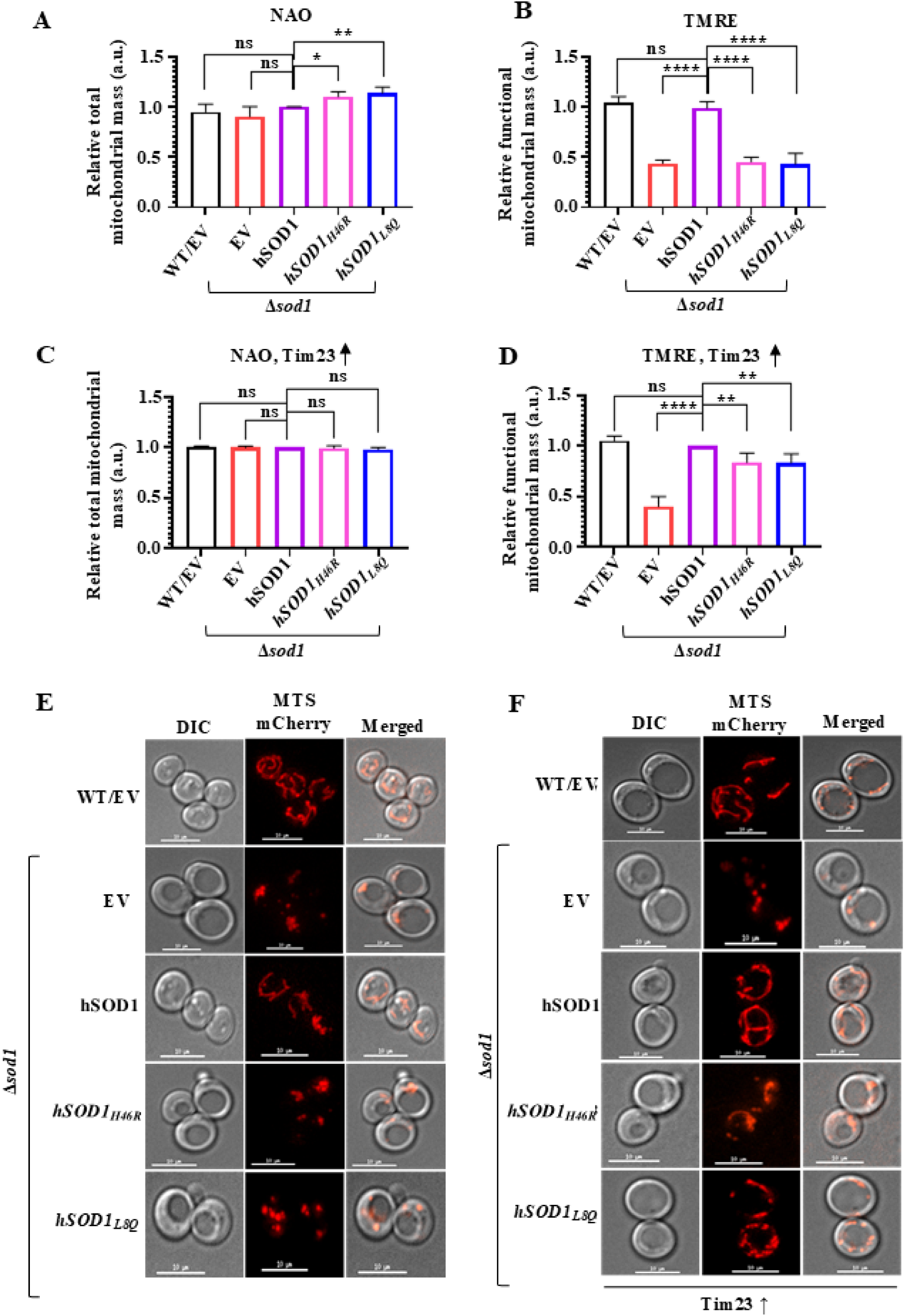
Determination of mitochondrial health parameters uponTim23 overexpression. (A,C) Total mitochondrial mass estimation. An equal number of cells from the mid-log phase were incubated with 10 µM NAO, without **(A)** or with Tim23 overexpression **(C)**. The cells were analyzed by flow cytometry and represented as relative total mitochondrial mass compared to hSOD1. **(B,D)** Functional mitochondrial mass assessment. Yeast cells from the above-mentioned strains without **(B)** or with overexpression of Tim23 **(D)** were treated with 8.75 µM TMRE for 20 min at 30℃, followed by flow cytometry analysis. The statistical analysis was performed using one-way ANOVA with Fisher’s LSD multiple comparisons test for **≥** 3 independent experiments. The asterisks indicate p values, **, *p*< 0.01 ***; *p*< 0.001; ****, *p*< 0.0001). **(E,F)** Mitochondrial morphology determination. Yeast cells expressing MTS-mCherry were analyzed by fluorescence microscopy to assess mitochondrial morphology under EV and Tim23 overexpressed **(F)** conditions. Scale bar: 10 µm.

Next, we overexpressed Tim23 in WT and mutant backgrounds and measured both total and functional mitochondrial mass. The Δ*sod1* cells showed reduced inner membrane potential even after Tim23 overexpression, as well as total mitochondrial content similar to hSOD1 (**Fig 8C-D**). In contrast, Tim23 overexpression reversed the increase in total mitochondrial mass observed in hSOD1 mutants (**Fig 8C**). Furthermore, Tim23 overexpression partially restored mitochondrial inner membrane potential in *hSOD1_H46R_* and *hSOD1_L8Q_*, indicating a partial rescue of dysfunctional mitochondria in these toxic hSOD1 mutants (**Fig 8C-D**). As a second parameter for mitochondrial integrity, we analyzed organellar morphology using fluorescence microscopy. WT and hSOD1 displayed a reticular morphology, whereas Δ*sod1* and hSOD1 mutants exhibited fragmented or punctate mitochondrial forms (**Fig 8E**, **S9A Fig**). Strikingly, overexpressing Tim23 partially rescued mitochondrial morphology in *hSOD1_H46R_* and *hSOD1_L8Q_,* consistent with recovery of ETC complex activity, inner membrane potential, and cellular growth (**Fig 8F**, **S9B Fig**). However, Tim23 overexpression had little or no effect on the punctate mitochondrial morphology of Δ*sod1* cells, which correlated with the growth pattern seen under overexpression conditions (**Fig 8E,F**). Tim23 overexpression did not alter mitochondrial morphology in WT+EV or hSOD1 strains (**Fig 8E,F**, **S9B Fig**). Altogether, Tim23 overexpression can partially restore mitochondrial integrity in the ALS-causing hSOD1 mutants.

**Fig 9.**
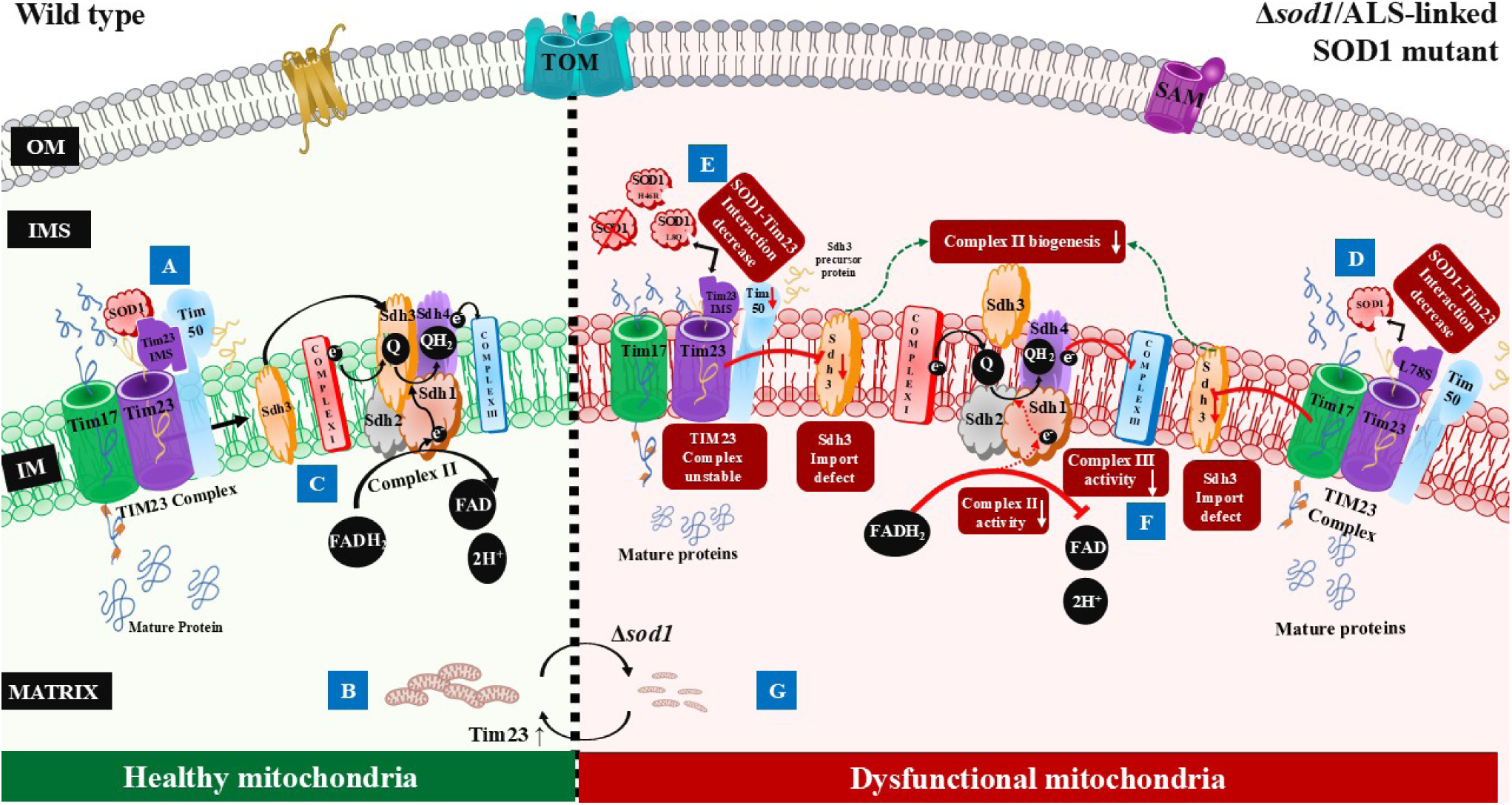
Model depicting the novel cross-talk between SOD1 and Tim23 modulating Sdh3 import to maintain mitochondrial health. **(A)** In healthy cells *(left panel)*, IMS-localized SOD1 dynamically interacts with Tim23, and SOD1-Tim23 interaction maintains TIM23^CORE^ complex stability and Sdh3 import. **(B)** Mitochondrial protein import is crucial for maintaining mitochondrial homeostasis. **(C)** Normal Sdh3 levels are crucial for ETC complex II activity. **(D)** In Tim23 mutant cells (*right panel*), the Sdh3 level is modulated by a reduced interaction of SOD1-Tim23 in the *tim23_L78_*_S_ mutant. **(E)** In cells lacking *SOD1* or expressing ALS-linked SOD1 mutants, Sdh3 import is defective due to decreased Tim23 and Tim50 protein levels and Tim23^CORE^ complex stability. **(F)** Reduced levels of Tim23 and Tim50 resulted in reduced activity of ETC complex II, leading to mitochondrial dysfunction. **(G)** In ALS-linked SOD1 mutants, fragmented mitochondria and reduced functional mitochondrial mass dictate disrupted mitochondrial integrity. IM-Mitochondrial inner membrane, (↓) - decrease.

## Discussion

ALS is an adult-onset neurodegenerative disease that selectively causes the death of motor neurons of the brain and spinal cord. The discovery of SOD1 mutations in ALS was a significant breakthrough in understanding disease pathogenesis. However, even after 30 years, the mechanism of cytotoxicity in ALS remains unknown. A mutation in SOD1 results in oligomerization of the protein, and a fraction of these misfolded species tends to accumulate in mitochondria, leading to organellar dysfunction. Although it is known that SOD1 mutants affect several mitochondrial functions, the molecular mechanism by which ∼5% of the total cellular SOD1 fraction residing in the organelle impairs various processes remains enigmatic.

In the present study, we propose a dynamic association between SOD1 mutants and mitochondrial IM pre-sequence translocase machinery. Under normal physiological conditions, SOD1 interacts with the TIM23 complex to stabilize the Tim23-Tim17-Tim50-Tim21 (23-17-50-21), Tim23-17-50, and Tim23-Tim17 complexes. The residues (L78) of intrinsically disordered IMS domain of Tim23 could play a crucial role in this interaction and govern the import of Tim23-specific substrate protein Sdh3. This results in effective ETC complex activity, preserving mitochondrial membrane potential and morphology. However, in hSOD1 mutants, binding is abrogated, resulting in decreased Sdh3 translocation and mis regulation of Complex II activity, leading to mitochondrial dysfunction. Strikingly, Tim23 overexpression partially rescued ETC complex activity and mitochondrial integrity. Hence, Tim23 overexpression partly restores cellular growth defects caused by the toxic mutant hSOD1.

The observed link between SOD1 and the TIM23 complex is not limited to diseased conditions. A conserved interaction between the pre-sequence translocase machinery and SOD1 was detected, even under physiological conditions, suggesting a novel function for SOD1 as a signaling molecule in mitochondrial biogenesis. Deleting SOD1 abolished its interaction with the TIM23 complex, thereby affecting mitochondrial function. Reduced SOD1 expression is observed in several other diseases, including Alzheimer’s disease and diabetic nephropathy [42–44]. Moreover, the steady-state levels of Tim23 were decreased in the substantia nigra of Parkinson’s disease, and its overexpression partially protected against neuronal death [45]. Mitochondrial dysfunction is a hallmark of these conditions, and our study may provide a basis for analyzing the functionality of the SOD1-TIM23 complex in diseases beyond ALS.

The ALS-linked mutations in hSOD1 were found throughout the protein, with more than 160 reported in the 154-amino-acid protein. In the present study, structurally and functionally distant SOD1 mutants were selected to understand the overall pathomechanism of ALS. The *hSOD1_H46R_* is an unusual mutant at the copper-binding site that results in the loss of dismutase activity but a slow progression of fALS [46, 47]. In contrast, *hSOD1_L8Q_* is located at the dimer interface, and the mutation results in rapid disease progression, with a lifespan of about 8 months from onset [48]. Even though the mutants have different toxicity levels, they showed a similar effect on the TIM23 complex association and mitochondrial function. Therefore, it is reasonable to believe that the mechanism of mitochondrial dysfunction reported here might be universal for most of the mutants of SOD1.

The mitochondrial dysfunction observed is mainly due to altered ETC complex activity in mutant hSOD1. These findings are further supported by previous reports suggesting altered mitochondrial bioenergetics. Several reports also predicted differential activities of ETC complex components, which hampered overall ATP production [21, 49]. However, to date, the exact mechanism and molecular signatures underlying the impaired ETC complex functions modulated by mutant hSOD1 remain poorly understood. Here, for the first time, the downregulation of Sdh3, a part of Complex II, was detected as a probable factor in initiating altered ETC activity. Decreased Sdh3 expression reduced Complex II activity, thereby affecting overall ETC function, as evidenced by Complex II+III activity. Therefore, the SOD1 interaction with the TIM23 complex may act as a signal for the Sdh3 expression, which, when impaired, alters ETC complex activity. The role of SOD1 in redox signalling is well established, with a significant portion of cellular SOD1 sensing nutrient availability to regulate energy metabolism [50, 51]. Sdh3 primarily functions in the mitochondrial TCA cycle and serves as a link between carbon metabolism and oxidative phosphorylation [52, 53]. The regulation of Sdh3 by the SOD1-TIM23 complex interaction may offer an exciting avenue for further study of SOD1 redox signalling. ALS motor neurons exhibit a metabolic switch in which the cells depend on glycolysis rather than oxidative phosphorylation for energy production [54, 55]. In line with this, decreased Sdh3 expression may provide a mechanism used by the mutants in pathogenic conditions.

The overexpression of Tim23 partially rescued the mitochondrial dysfunction and cellular growth in hSOD1 mutants. The effect of overexpression was mainly on the recovery of ETC complex activity, thereby restoring mitochondrial function. The results suggest that Tim23 overexpression increases the mitochondrial pool of the substrate protein Sdh3, which may explain the improved Complex II activity and mitochondrial integrity. Further analysis is required to understand the underlying mechanism by which the TIM23 complex mediates SOD1-induced downregulation of Sdh3. Also, evaluating other substrates of the TIM23 complex may provide a more detailed explanation of the growth rescue observed under Tim23 overexpression.

In conclusion, the study predicts a novel regulation of mitochondrial biogenesis by presequence translocase machinery in ALS-linked SOD1 mutants. Identifying the SOD1-TIM23 complex interaction governing Sdh3 translocation strengthens the possible role of SOD1 in metabolic signalling. Decreased SOD1-TIM23 complex interaction in ALS-associated mutants led to defective ETC function and impaired mitochondrial function, shown by Sdh3 downregulation. This highlights a plausible mechanism for altered ETC complex activity, featuring a specific molecule regulated by mutant SOD1 *via* presequence translocase machinery. Interestingly, overexpression of Tim23 partially rescued cellular growth defects caused by mutant SOD1 by suppressing mitochondrial dysfunction. Therefore, the results open avenues for exploring the role of the TIM23 complex in the design of therapeutic ALS strategies.

## Materials and Methods

### Yeast strains, cloning, and cell culture media

The *in vivo* studies were performed using haploid W303 background strain of *S. cerevisiae* (Genotype: *ade2-1/ade2-1 his3-11/his3-11, 15 ura3-1/15 ura3-1 leu2-3/leu2-3, 112 trp1-1/112 trp1-1, can1-100/can1-100 GAL2/GAL2, met2-1/met2-1 lys2-2/lys2-2* and W303 derivative, PJ53 background strain of *S. cerevisiae* (Genotype: *trp1-1/trp1-1, ura3-1/ura3-1, leu2-3,112/leu2-3/112, his3-11,15/his3-11,15 ade2-1/ade2-1, can1-100/can1-100 GAL2+/GAL2+, met2-Δ1/met2-Δ1, lys2-Δ2/lys2-Δ2*). The genomic copy of *SOD1* was deleted by homologous recombination using a previously published protocol, where *SOD1* was replaced by the hygromycin-resistant (*Hph*) gene from the pYM16 vector using primers P1 and P2 mentioned in the list of primers [56]. The transformants obtained on YPD (1% yeast extract, 2% peptone, and 2% dextrose) supplemented with Hygromycin (280 µg/ml) were further confirmed by the colony PCR analysis. *SOD1* was amplified from the genomic DNA of the PJ53 strain, followed by cloning with the C-terminus FLAG tag in the yeast expression vector pRS416GPD (Gifted by Prof. Elizabeth A. Craig’s laboratory, University of Wisconsin-Madison). To generate Tim23 mutant strains, we used the haploid Δ*tim23* knockout strain in the W303 background (kindly provided by the laboratory of Prof. Elizabeth A. Craig, University of Wisconsin-Madison). Tim23 mutants were cloned into the yeast expression vector pRS314 with the native promoter and confirmed by sequencing. The mutants encoding plasmids were transformed into the Δ*tim23* haploid strain harbouring a functional copy of the *TIM23* gene on a URA3-based plasmid. The presence of the *URA3* gene allowed for the selection of cells that had lost the URA-based plasmid carrying the wild-type *TIM23* gene on the 5-fluoro uracil (5-FOA) His-Trp-media incubated at 30°C.

hSOD1 and its mutants were cloned into the yeast expression vector pRS416 under the *GPD* promoter and confirmed by sequencing. For overexpression of Tim23 and Dic1, the respective genes were cloned into the pRS414 vector under the *TEF* promoter. Yeast cells were cultured in synthetic complete dextrose (SCD) medium containing 2% dextrose, 0.67% yeast nitrogen base, and 0.67% amino acid supplements, with uracil (Ura-), leucine (Leu-), and tryptophan (Trp-) omitted as required. GST-tagged *SOD1* was generated by cloning the *SOD1* gene into the C-terminus of GST in the pGEX-KG vector and confirmed by sequencing.

HEK293T cells were grown in Advanced DMEM/F-12 (Dulbecco’s Modified Eagle Medium/Ham’s F-12) (Gibco^TM^) supplemented with 10% fetal bovine serum (FBS, GibcoFFSDDFTM), 1% Glutamax (Gibco^TM^), and 1% penicillin-streptomycin (Gibco^TM^).

### Antibody profiles

Antibodies against FLAG (1:1000 dilution, F1804, Sigma), hSOD1 (1:3000 dilution, HPA001401, Sigma), and Por1 (1:5000 dilution, 16G9E6BC4, Thermo Fisher) were purchased from the manufacturer. Antibodies specific to Tim54, Tim22, Tim18, Sdh3, and Tim10 have been described previously [57]. Briefly, the antibody against Tim54 (1:1000 dilution) was a gift from Prof. Toshiya Endo (Kyoto Sangyo University, Japan). Antisera against Tim22 (1:250 dilution) and Dic1 (1:250 dilution) were gifted by Prof. Agnieszka Chacinska (Warsaw University, Poland). The Sdh3 (1:250 dilution) antibody was a gift from Prof. Nikolaus Pfanner (University of Freiburg, Germany). The antibody against Tim10 was a gift from Prof. Carla M. Koehler (UCLA, USA). Antibodies specific to yeast proteins, Tim44 (1:3000 dilution), Hsp60 (1:3000 dilution), Mdj1 (1:3000 dilution), and Ydj1 (1:5000 dilution), were kindly gifted by Prof. Elizabeth A. Craig’s laboratory (University of Wisconsin-Madison). Tim18 (1:2500 dilution) antibody was raised without the presequence in rabbits by Abgenex Pvt. Ltd. Anti-Tim23 (1:3000 dilution) and anti-Tim50 (1:3000 dilution) were generated against the N-terminus (1-98) and C-terminus (133-476) amino acids, respectively, by the Abgenex Pvt. Ltd. Tim21 (1:3000 dilution) and Tim17 (1:250 dilution) antibodies were raised in rabbits by Abgenex Pvt. Ltd. Antibodies against the respiratory complex proteins are described previously [58]. Human proteins specific antibodies are specified in the previously published report [59].

### *In vivo* precursor accumulation analysis

Wild-type and Tim23 mutant yeast strains were grown at a permissible temperature to early log phase, then exposed to heat shock at 37°C for 10h to induce the phenotype. For Tim23 mutant strains with a *SOD1* deletion background, heat shock was given at 34°C for 8h. Samples were analyzed by SDS-PAGE and immunoblotting using Mdj1 and Hsp60 antibodies to assess precursor accumulation, with Ydj1 as a cytosolic loading control [58].

### Blue native polyacrylamide gel electrophoresis (BN-PAGE)

BN-PAGE was performed as previously described [60]. Briefly, mitochondria (200 µg/ml) were mixed with 100 μl of a digitonin buffer (1% digitonin, 50 mM NaCl, 50 mM imidazole, 2 mM 6-aminocaproic acid, 1 mM EDTA, pH 7.0) and kept for 1 hour at 4°C. The insoluble part was separated by spinning at 16,900 g for 20 minutes at 4°C. Next, 15 μl of sample buffer (containing 2.5 μl of 5% Coomassie stain and 12.5 μl of 50% glycerol) was added to 100 μl of the clear fraction. The samples were loaded onto a 6–16% gradient gel, followed by western blot and detection with antibodies specific to Tim22 and Tim23.

### Quantification of protein half-lives

Wild-type and mutant yeast cells were grown in liquid Ura-dropout media to mid-log phase at 30°C. 50 µg/ml of cycloheximide (Sigma) was added to the culture to inhibit protein translation. Following the treatment, an equal number of cells were collected at 0, 4, 8, and 12 h. Cell lysates were prepared and separated on SDS-polyacrylamide gels, and the proteins were immunoblotted with antibodies against Tim23, Tim50, FLAG, and Ydj1.

### Co-IP analysis

Mitochondria from yeast and human HEK293T cells were prepared as previously described [61,62]. 5 mg of mitochondria were broken open in a solution containing 25 mM Tris-HCl (pH 7.5), 10% glycerol, 80 mM KCl, 5 mM EDTA, 1 mM PMSF, and 1% digitonin. Unbroken mitochondria were removed by spinning at 14000 rpm for 20 min at 4℃. The clear liquid was mixed with beads attached to antibodies for 12h at 4℃: protein G Sepharose beads for FLAG or protein A Sepharose beads for Tim23. Unbound proteins were washed away with a 0.5% digitonin solution. The remaining proteins were separated by SDS-PAGE and detected by immunoblotting with specific antibodies for yeast or human samples.

### Protein purification and GST pull-down assay

*SOD1* cloned into the pGEX-KG vector was transformed into an *Escherichia coli* (RIL) expression strain. The proteins were induced at 30℃ for 6h, followed by purification, as reported earlier, with a minor modification [63]. Tris-HCl pH 7.5 was used instead of phosphate buffer in the purification protocol. For GST pull-down analysis, 5 mg of mitochondria from the WT strain were lysed in the pull-down buffer (50 mM Tris-HCl, pH 7.5, 100 mM NaCl, 10% Glycerol, and 1 mM PMSF) with 0.5% NP-40. After centrifugation, the clear lysate was incubated with 5 µM of purified protein attached to the beads for 12h at 4℃. Samples were washed twice with pull-down buffer, separated by SDS-PAGE, and detected by Coomassie staining or specific antibodies.

### ETC complex activity

ETC complex II and II+III activity was performed as described earlier [64]. Briefly, for complex II activity, 30 µg of mitochondria were incubated for 1 min at 30℃ in phosphate buffer (0.5 M, pH 7.5), fatty acid-free BSA (1 mg/ml, Sigma), and succinate (20 µg/ml, Sigma). To check the specific activity, complexes I, III, and IV were inhibited by the addition of Rotenone (2 µg/ml, Sigma), Antimycin (10 µg/ml, Sigma), and sodium azide (2 mM, Sigma) for 3 min at 30℃. Decylubiquinone (0.01 mM, Sigma) was added to the reaction mixture, followed by incubation for 60 sec at 30℃. The reaction was initiated by adding 2,6-dichlorophenolindophenol sodium salt hydrate (DCPIP, Sigma), and the decrease in absorbance at 600 nm was monitored for 5 min. The complex II+III activity was performed in the presence of sodium azide. Briefly, 30 µg of yeast mitochondria were incubated in the phosphate buffer (0.5 M, pH 7.5) and succinate (20 µg/ml, Sigma) for 60 sec at 30℃. The reaction mixture was incubated with sodium azide (2 mM, Sigma) for 3 min at 30℃, followed by the addition of 1 mM oxidized cytochrome C (Sigma). The increase in absorbance was recorded at 550 nm for 5 min.

### Flow cytometry analysis

We measured total mitochondrial mass using Nonyl Acridine Orange (NAO, Invitrogen). Yeast cells (*A_600_* = 0.1) grown at 34℃, followed by incubation with 10 µM NAO at 30℃ for 20 min while shaking at 900 rpm. After washing the cells with a salt solution (PBS), we measured the results using a machine that detects light at 496 nm (excitation) and 519 nm (emission). We also measured mitochondrial energy potential using the dye TMRE (8.75 µM at 30℃ for 30 min, Invitrogen) and detected light at 552 nm (excitation) and 574 nm (emission).

### Fluorescence microscopy

For mitochondrial morphology visualization, cells with EV or Tim23 overexpression were grown till the mid-log phase. Mitochondria were specifically decorated with MTS-mCherry plasmid as described earlier [38]. Cells were harvested by centrifugation at 5000 rpm, 5 min at room temperature, followed by resuspension in PBS. Cells were mounted on 2% agarose pads and imaged with a DeltaVision Elite fluorescence microscope (GE Healthcare) using a 100× objective lens and constant exposure. The excitation and emission wavelengths of 587 nm and 610 nm were used for mCherry. The images were deconvolved and examined using SoftWoRx 6.1.3 software (GE Healthcare) and ImageJ software.

### Statistical analysis

Statistical analyses of biological replicates were performed using one-way ANOVA with Fisher’s LSD multiple comparisons test and unpaired t-tests, comparing all columns in the specified samples. Statistical significance (p-value) is indicated by asterisks as follows: \**p*< 0.05, \*\**p* < 0.01, \*\*\**p*< 0.001, \*\*\*\**p*< 0.0001.

## List of primers used in the study

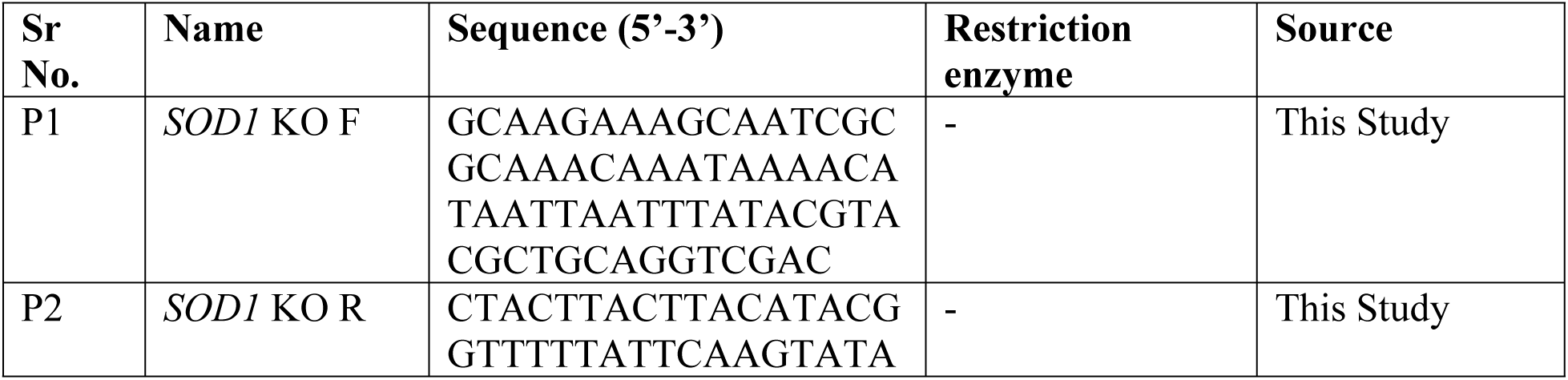
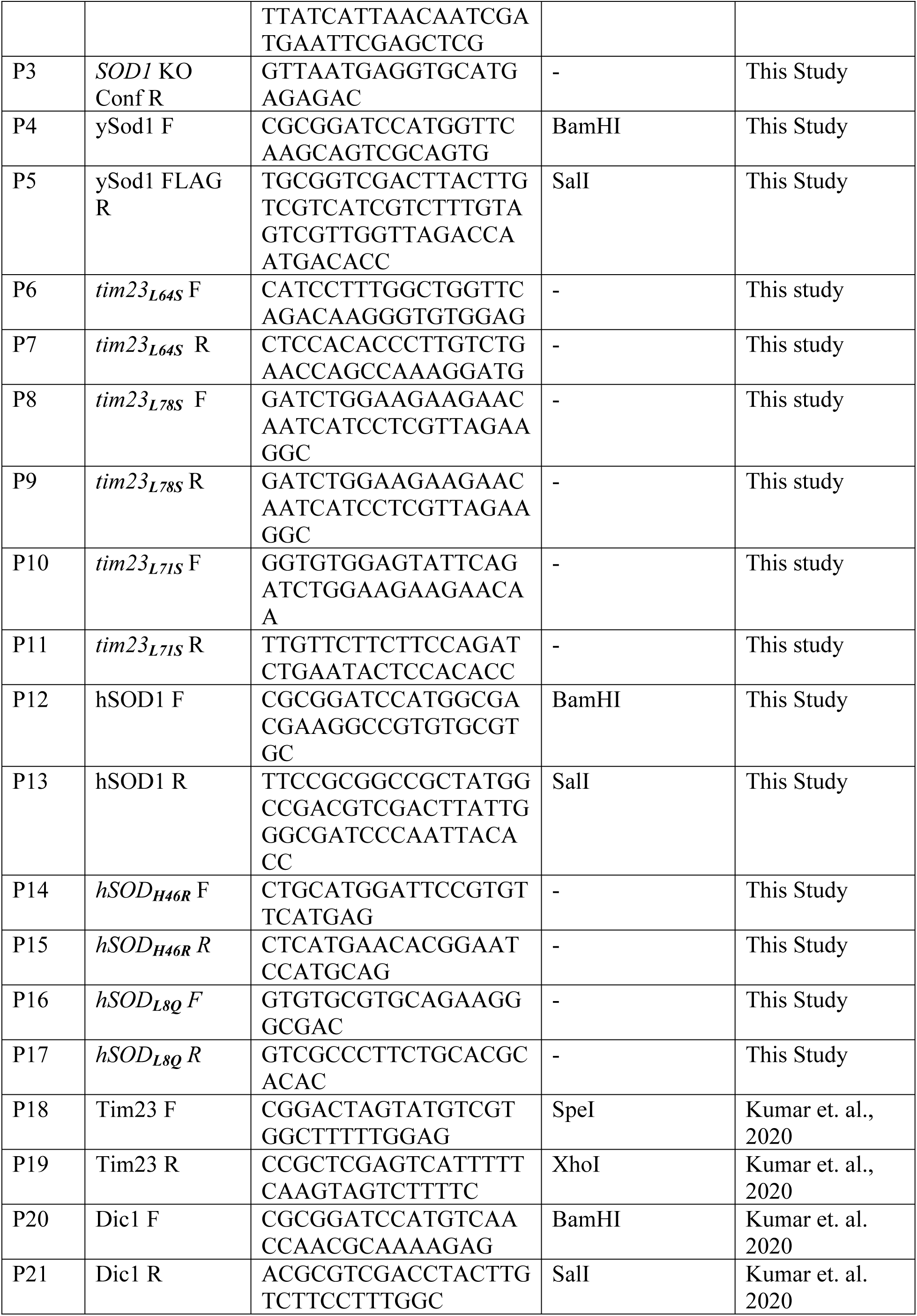

## List of strains used in the study

| Sr. No. | Name | Genotype | Source |
| --- | --- | --- | --- |
| 1 | WT | *trp1-1/trp1-1, ura3-1/ura3-1, leu2-3,112/leu2-3/112, his3-11,15/his3-11,15 ade2-1/ade2-1, can1-100/can1-100 GAL2+/GAL2+, met2-Δ1/met2-Δ1, lys2-Δ2/lys2-Δ2* | Sinha et. al., 2010 |
| 2 | *Δsod1* | *trp1-1/trp1-1, ura3-1/ura3-1, leu2-3,112/leu2-3/112, his3-11,15/his3-11,15 ade2-1/ade2-1, can1-100/can1-100 GAL2+/GAL2+, met2-Δ1/met2-Δ1, lys2-Δ2/lys2-Δ2, sod1Δ::CgHph* | This Study |
| 3 | *Δsod1/ySOD1-FLAG* | *trp1-1/trp1-1, ura3-1/ura3-1, leu2-3,112/leu2-3/112, his3-11,15/his3-11,15 ade2-1/ade2-1, can1-100/can1-100 GAL2+/GAL2+, met2-Δ1/met2-Δ1, lys2-Δ2/lys2-Δ2, sod1Δ::CgHph [pRS416GPD-ysod1FLAG]* | This Study |
| 4 | *Δtim23* | *trp1-1/trp1-1, ura3-1/ura3-1, leu2-3,112/leu2-3/112, his3-11,15/his3-11,15 ade2-1/ade2-1, can1-100/can1-100 GAL2+/GAL2+, met2-Δ1/met2-Δ1, lys2-Δ2/lys2-Δ2, Tim23Δ::CgHIS3 [pRS316-Tim23]* | This study |
| 5 | *Δtim23/tim23<sub>L64S</sub>* | *trp1-1/trp1-1, ura3-1/ura3-1, leu2-3,112/leu2-3/112, his3-11,15/his3-11,15 ade2-1/ade2-1, can1-100/can1-100 GAL2+/GAL2+, met2-Δ1/met2-Δ1, lys2-Δ2/lys2-Δ2, Tim23Δ::CgHIS3 [pRS314-Tim23<sub>L64S</sub>]* | This study |
| 6 | *Δtim23/tim23<sub>L78S</sub>* | *trp1-1/trp1-1, ura3-1/ura3-1, leu2-3,112/leu2-3/112, his3-11,15/his3-11,15 ade2-1/ade2-1, can1-100/can1-100 GAL2+/GAL2+, met2-Δ1/met2-Δ1, lys2-Δ2/lys2-Δ2, Tim23Δ::CgHIS3 [pRS314-Tim23<sub>L78S</sub>]* | This study |
| 7 | *Δtim23/tim23<sub>L71S</sub>* | *trp1-1/trp1-1, ura3-1/ura3-1, leu2-3,112/leu2-3/112, his3-11,15/his3-11,15 ade2-1/ade2-1, can1-100/can1-100 GAL2+/GAL2+, met2-* | This study |
|  |  | *Δ1/met2-Δ1, lys2-Δ2/lys2-Δ2, Tim23Δ::CgHIS3 [pRS314-Tim23<sub>L71S</sub>]* |  |
| 8 | *Δsod1/Δtim23:Tim23<sub>314</sub>* | *trp1-1/trp1-1, ura3-1/ura3-1, leu2-3,112/leu2-3/112, his3-11,15/his3-11,15 ade2-1/ade2-1, can1-100/can1-100 GAL2<sup>+</sup>/GAL2<sup>+</sup>, met2-Δ1/met2-Δ1, lys2-Δ2/lys2-Δ2, sod1Δ::CgHph, Tim23Δ::CgHIS3 [pRS314-Tim23]* | This study |
| 9 | *Δsod1/Δtim23:tim23<sub>L64S</sub>* | *trp1-1/trp1-1, ura3-1/ura3-1, leu2-3,112/leu2-3/112, his3-11,15/his3-11,15 ade2-1/ade2-1, can1-100/can1-100 GAL2<sup>+</sup>/GAL2<sup>+</sup>, met2-Δ1/met2-Δ1, lys2-Δ2/lys2-Δ2, sod1Δ::CgHph, Tim23Δ::CgHIS3 [pRS314-Tim23<sub>L64S</sub>]* | This study |
| 10 | *Δsod1/Δtim23:tim23<sub>L78S</sub>* | *trp1-1/trp1-1, ura3-1/ura3-1, leu2-3,112/leu2-3/112, his3-11,15/his3-11,15 ade2-1/ade2-1, can1-100/can1-100 GAL2<sup>+</sup>/GAL2<sup>+</sup>, met2-Δ1/met2-Δ1, lys2-Δ2/lys2-Δ2, sod1Δ::CgHph, Tim23Δ::CgHIS3 [pRS314-Tim23<sub>L78S</sub>]* | This study |
| 11 | *Δsod1/Δtim23:tim23<sub>L71S</sub>* | *trp1-1/trp1-1, ura3-1/ura3-1, leu2-3,112/leu2-3/112, his3-11,15/his3-11,15 ade2-1/ade2-1, can1-100/can1-100 GAL2<sup>+</sup>/GAL2<sup>+</sup>, met2-Δ1/met2-Δ1, lys2-Δ2/lys2-Δ2, sod1Δ::CgHph, Tim23Δ::CgHIS3 [pRS314-Tim23<sub>L78S</sub>]* | This study |
| 12 | hSOD1 | *trp1-1/trp1-1, ura3-1/ura3-1, leu2-3,112/leu2-3/112, his3-11,15/his3-11,15 ade2-1/ade2-1, can1-100/can1-100 GAL2<sup>+</sup>/GAL2<sup>+</sup>, met2-Δ1/met2-Δ1, lys2-Δ2/lys2-Δ2, sod1Δ::CgHph [pRS416GPD-*hsod1*]* | This Study |
| 13 | *Δsod1/hSOD1<sub>H46R</sub>* | *trp1-1/trp1-1, ura3-1/ura3-1, leu2-3,112/leu2-3/112, his3-11,15/his3-11,15 ade2-1/ade2-1, can1-100/can1-100 GAL2<sup>+</sup>/GAL2<sup>+</sup>, met2-Δ1/met2-Δ1, lys2-Δ2/lys2-Δ2, sod1Δ::CgHph [pRS416GPD-*hsod1H46R*]* | This Study |
| 14 | Δsod1/hSOD<sub>L8Q</sub> | *trp1-1/trp1-1, ura3-1/ura3-1, leu2-3,112/leu2-3/112, his3-11,15/his3-11,15 ade2-1/ade2-1, can1-100/can1-100 GAL2+/GAL2+, met2-Δ1/met2-Δ1, lys2-Δ2/lys2-Δ2, sod1Δ::CgHph [pRS416GPD-*hsod1L8Q*]* | This Study |

## List of Reagents used in the study

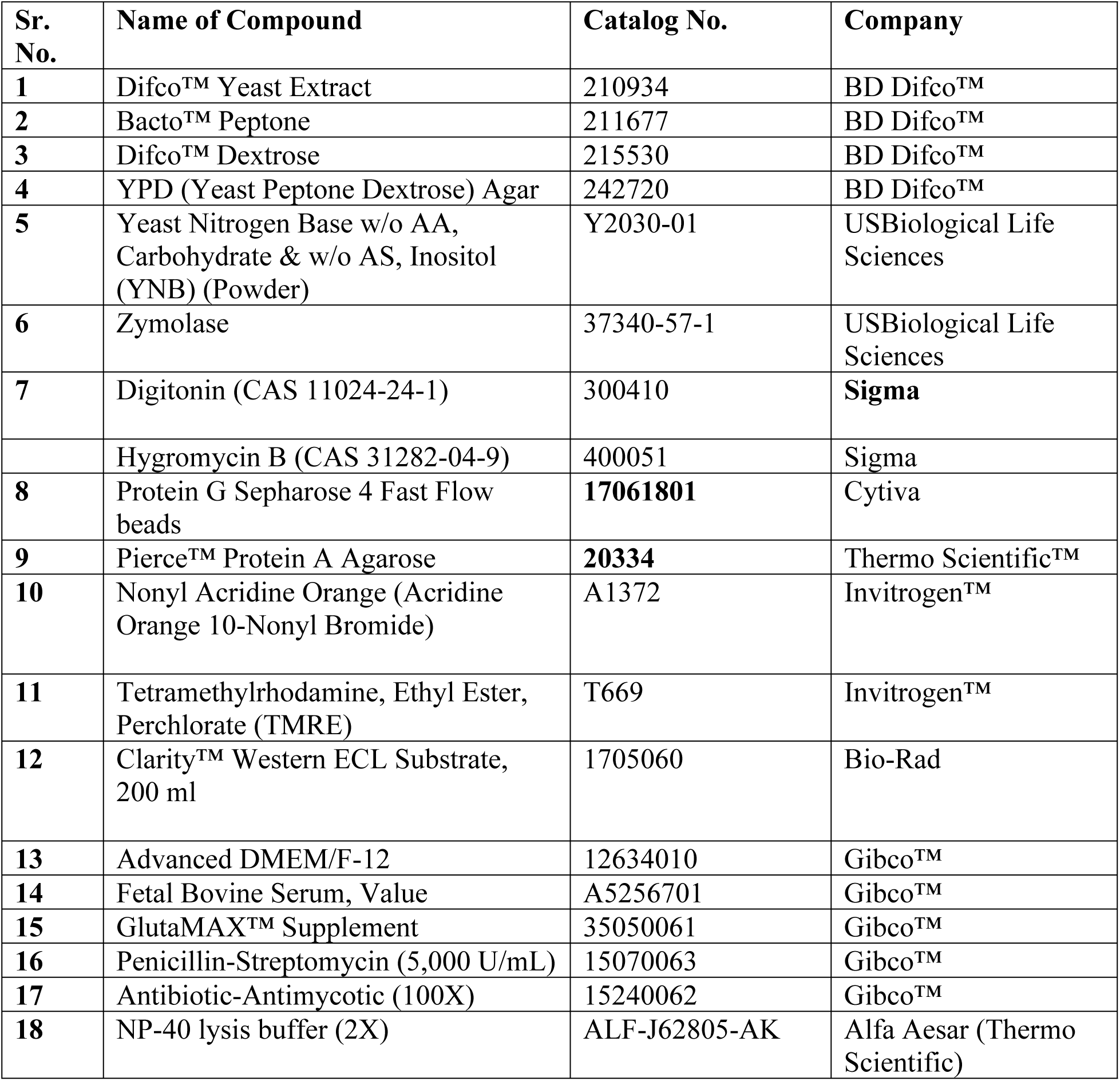

| Sr. No. | Name of Compound | Catalog No. | Company |
| --- | --- | --- | --- |
| 1 | Difco™ Yeast Extract | 210934 | BD Difco™ |
| 2 | Bacto™ Peptone | 211677 | BD Difco™ |
| 3 | Difco™ Dextrose | 215530 | BD Difco™ |
| 4 | YPD (Yeast Peptone Dextrose) Agar | 242720 | BD Difco™ |
| 5 | Yeast Nitrogen Base w/o AA, Carbohydrate & w/o AS, Inositol (YNB) (Powder) | Y2030-01 | USBiological Life Sciences |
| 6 | Zymolase | 37340-57-1 | USBiological Life Sciences |
| 7 | Digitonin (CAS 11024-24-1) | 300410 | **Sigma** |
|  | Hygromycin B (CAS 31282-04-9) | 400051 | Sigma |
| 8 | Protein G Sepharose 4 Fast Flow beads | **17061801** | Cytiva |
| 9 | Pierce™ Protein A Agarose | **20334** | Thermo Scientific™ |
| 10 | Nonyl Acridine Orange (Acridine Orange 10-Nonyl Bromide) | A1372 | Invitrogen™ |
| 11 | Tetramethylrhodamine, Ethyl Ester, Perchlorate (TMRE) | T669 | Invitrogen™ |
| 12 | Clarity™ Western ECL Substrate, 200 ml | 1705060 | Bio-Rad |
| 13 | Advanced DMEM/F-12 | 12634010 | Gibco™ |
| 14 | Fetal Bovine Serum, Value | A5256701 | Gibco™ |
| 15 | GlutaMAX™ Supplement | 35050061 | Gibco™ |
| 16 | Penicillin-Streptomycin (5,000 U/mL) | 15070063 | Gibco™ |
| 17 | Antibiotic-Antimycotic (100X) | 15240062 | Gibco™ |
| 18 | NP-40 lysis buffer (2X) | ALF-J62805-AK | Alfa Aesar (Thermo Scientific) |

## Acknowledgments

We thank **Prof. Elizabeth A. Craig** for yeast strains, vectors, and yeast-specific antibodies. We thank **Prof. Toshiya Endo** (Kyoto Sangyo University, Japan), Prof. Agnieszka Chacinska (Warsaw University, Poland), Prof. Nikolaus Pfanner (University of Freiburg, Germany), and **Prof. Carla M. Koehler** (UCLA, USA) for the antibodies. We are grateful to Vinaya Vishwanathan and Abhishek Kumar for providing critical insights into the experiments and vector constructs. We thank the Department of Biochemistry’s central facility and the FACS facility for providing the required instruments and technicians. **Patrick D’Silva** acknowledges funding from ANRF and DST-FIST for this study. **Tejashree Pradip Waingankar** acknowledges CSIR for this study. **Aakansha Paliwal** acknowledges PMRF for this study. **Anjali Deep** acknowledges IISC fellowship.

## Author contributions

**Conceptualization:** Tejashree Pradip Waingankar, Patrick D’Silva.

**Data curation:** Tejashree Pradip Waingankar, Aakansha Paliwal, Anjali Deep, and Patrick D’Silva.

**Formal analysis:** Tejashree Pradip Waingankar, Patrick D’Silva.

**Funding acquisition:** Patrick D’Silva.

**Investigation:** Tejashree Pradip Waingankar, Aakansha Paliwal, Anjali Deep.

**Methodology:** Tejashree Pradip Waingankar, Aakansha Paliwal, Anjali Deep.

**Project administration:** Patrick D’Silva.

**Resources:** Patrick D’Silva.

**Supervision:** Patrick D’Silva.

**Validation:** Tejashree Pradip Waingankar, Aakansha Paliwal, Patrick D’Silva.

**Visualization:** Tejashree Pradip Waingankar, Aakansha Paliwal, Anjali Deep, Patrick D’Silva.

**Writing:** Tejashree Pradip Waingankar, Aakansha Paliwal, Anjali Deep and Patrick D’Silva.

## Abbreviations

ALS: Amyotrophic Lateral Sclerosis
FDA: Food and Drug Administration
sALS: sporadic ALS
fALS: familial ALS
SOD1: Superoxide Dismutase 1
IMS: Inter membrane space
OM: Outer Membrane
IM: Inner Membrane
TOM: Translocase of OM
TIM: Translocase of IM
ETC: Electron Transport Chain
WT: Wild type
EV: Empty vector
Co-IP: Co-immunoprecipitation
Sdh3: Succinate dehydrogenase
NAO: 10-N-nonyl acridine orange
TMRE: Tetramethylrhodamine methyl ester
DCPIP: dichlorophenolindophenol sodium salt hydrate

